# Nitrogen-limitation independent control of *glnA* (glutamine synthetase) expression in *Escherichia coli* by urea, several amino acids, and post-transcriptional regulation

**DOI:** 10.1101/2023.01.17.524496

**Authors:** Karthik Urs, Philippe E. Zimmern, Larry Reitzer

**Author notes:** Address correspondence to Larry Reitzer. The University of Texas Dallas; Department of Biological Sciences. University of Texas Southwestern Medical Center; Department of Urology.

## Abstract

The expression of *glnA* (ammonia-assimilating glutamine synthetase) is high for uropathogenic *E. coli* grown in urine. Because *glnA* is part of an operon that codes for regulators of the nitrogen-regulated (Ntr) response, high *glnA* expression has been interpreted to suggest nitrogen limitation, which is unexpected because of the high urinary ammonia concentration and the extremely rapid bacterial growth. We present evidence that *glnA* expression does not result from nitrogen limitation. First, in the presence of ammonia, urea induced expression of *glnA* from the cAMP receptor protein (Crp)- dependent *glnAp1* promoter, which circumvents control from the nitrogen-regulated *glnAp2* promoter. This urea effect on *glnA* expression has not been previously described. Second, the most abundant amino acids in urine inhibited GS activity, based on reversal of the inhibition by glutamate and glutamine, and increased *glnA* expression. The relevance of these inhibitory amino acids in natural environments has not been previously demonstrated. Third, neither urea nor the inhibitory amino acids induced other Ntr genes, i.e., high *glnA* expression can be independent of other Ntr genes. Finally, the urea-dependent induction did not result in GlnA synthesis because of a previously undescribed translational control. We conclude that *glnA* expression in urea-containing environments does not imply growth rate-limiting nitrogen restriction and is consistent with rapid growth of uropathogenic *E. coli*.

**Significance:** Urinary tract infections (UTIs), often caused by *E. coli*, frequently become resistant to antibiotic treatment. Expressed metabolic genes during an infection could guide development of urgently-needed alternate or adjunct therapies. *glnA* (glutamine synthetase) is expressed during growth in urine, which implies growth-restricting nitrogen limitation. We show that *glnA* expression results from urinary amino acids that inhibit GlnA activity and urea, but not from nitrogen limitation. Urinary components will vary greatly between individuals which suggests corresponding variations in *glnA* expression. GlnA may be a metabolic vulnerability during UTIs, which may depend on a variable urinary composition. *glnA* expression may be important in a complex host-pathogen interaction, but may not be a good therapeutic target.

## Introduction

Urinary tract infections (UTIs) are common bacterial infections and UTI-associated *E. coli* (UTEC) causes most UTIs (1-3). During an infection, UTEC strains cycle between the bladder lumen and intracellularly within uroepithelial cells (4). Rapid growth occurs in both environments and is considered a virulence factor. Rapid growth is reflected in gene expression patterns, such as high expression of genes for the translational machinery (4-6). Antibiotic treatment is often effective but continued antibiotic use increases the risk of generating multidrug resistant organisms that can ultimately impact the outcome of other clinically important diseases that are treated with the same antibiotics (2, 7-9). Alternate or adjunct therapies are urgently needed (2, 4, 10), but potential targets of therapy have been elusive, in part because of an incomplete understanding of virulence. Likely targets include proteins that are expressed during an infection.

The genes for ammonia and iron acquisition are induced during growth in urine which implies multiple nutrient limitations that should be incompatible with rapid growth (11-13). Nitrogen limitation is implied by high *glnA* expression which is unexpected because urinary ammonia is high (> 20 mM) (13). The *glnA* product, glutamine synthetase (GS), catalyzes ATP-dependent glutamine synthesis from ammonia and glutamate. Multiple mechanisms control activity and synthesis of GS which is at the intersection of carbon and nitrogen metabolism (14). Figure 1 (black arrows) summarizes known regulation of GS activity and synthesis which includes direct inhibition of activity by several amino acids and nucleotides, a three-protein regulatory cascade that adenylylates and inactivates GS, transcriptional activation from the *glnAp1* promoter by carbon limitation, and transcriptional activation from the *glnAp2* promoter by nitrogen limitation. *glnA* is part of the *glnALG* operon that codes for regulators of a set of genes that respond to nitrogen limitation, which are collectively called Ntr (nitrogen-regulated) genes. Although not a direct regulatory factor, ammonia ― a precursor for glutamine synthesis ― suppresses *glnA* expression, except in urine.

**Figure 1.**
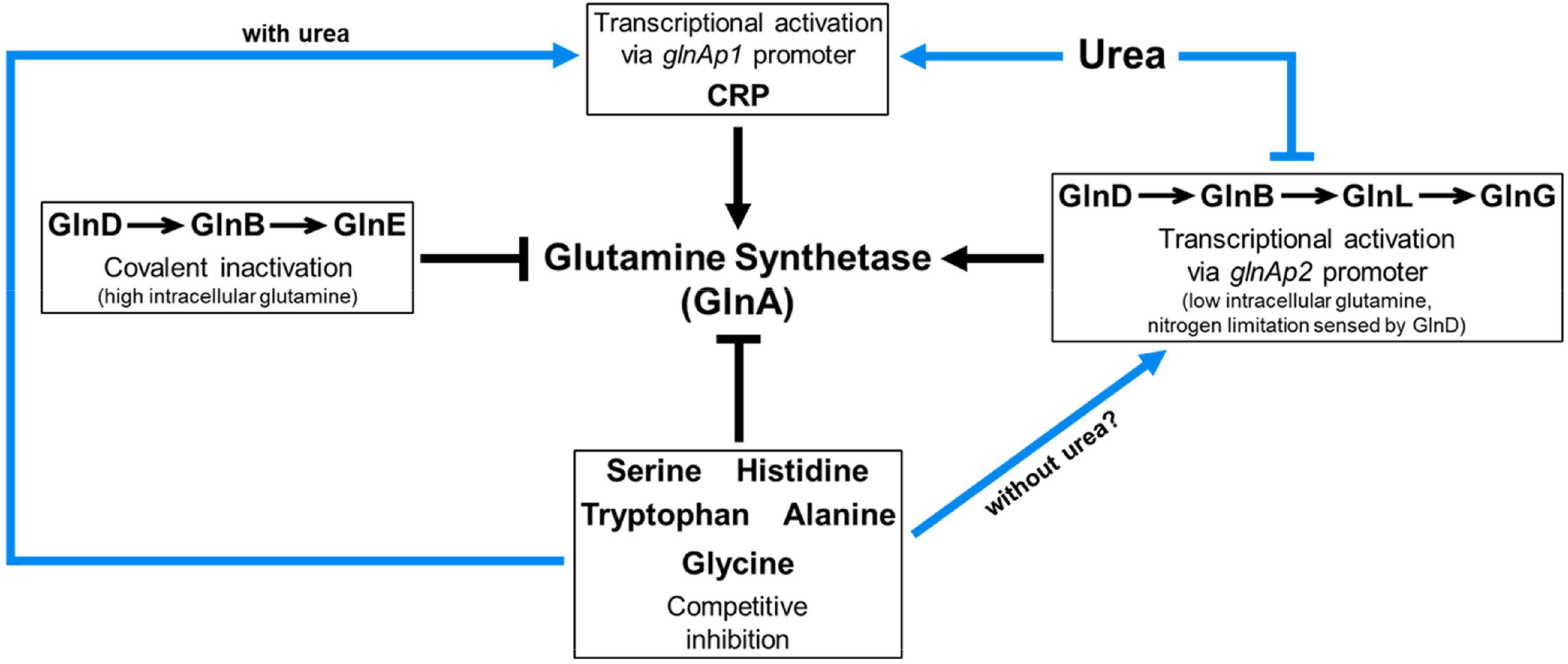
Overview of regulation of glutamine synthetase activity. Known factors that control GS activity and *glnA* expression are shown with black arrows, and factors that are identified in the present work are shown with blue arrows. First, high intracellular glutamine - mediated by the glutamine-sensing GlnD (uridylyltransferase/uridylyl-removing enzyme), and acting via GlnB (the regulatory protein PII), and GlnE (adenylyltransferase/deadenylylase) - covalently adenylylates and inactivates GS. Second, low intracellular glutamine - which is sensed by GlnD acting through GlnB, GlnL (a sensor kinase), and GlnG (a response regulator) - activates expression of *glnA* from the *glnAp2* promoter and a set of genes, which are collectively called Ntr (nitrogen-regulated) genes. Third, cyclic-AMP-Crp activates *glnA* expression from the *glnAp1* promoter. Fourth, urea increases *glnA* expression in ammonia-containing media from the *glnAp1* promoter, but impairs expression in nitrogen-limited media from the *glnAp2* promoter. Finally, several amino acids and nucleotides, including many that require glutamine for their synthesis, bind the glutamate- and nucleotide-binding sites and inhibit GS activity. In addition to the metabolites shown, glucosamine-6-phosphate, AMP and CTP inhibit activity. These inhibitory amino acids also affect *glnA* activity. Our results suggest that with urea *glnA* expression is initiated from the *glnAp1* promoter. Without urea, inconclusive evidence suggests expression initiated from the *glnAp2* promoter.

Our goal was to determine the factor(s) that could induce *glnA* expression in an ammonia-containing environment. We show that both urea and GS-inhibiting amino acids, which are abundant in urine, affect *glnA* expression (Fig. 1), and unexpectedly observe a previously undescribed translational control. We present the first description of (a) the effect of urea on *glnA* expression, and (b) the physiological relevance of amino acid inhibition of GS activity.

## Results

### Abundant urinary amino acids inhibit growth and GS activity

We grew UTI89 and W3110 in a minimal salts medium (MM) and a defined synthetic urine (SU) with basal components at urine-like concentrations. UTI89 and W3110 grew equally well in SU medium with glucose (Fig. 2), succinate (Fig. S1), and glycerol (Fig. S1) as the carbon source. Δ*glnA* derivatives were grown as negative controls. Doubling times were comparable between MM and SU media for UTI89 (Table S1), while the doubling time was slightly faster in MM-ammonia medium for W3110 (Table S2). (Tables S1 and S2 have the doubling time and final cell densities for all growth experiments).

**Figure 2:**
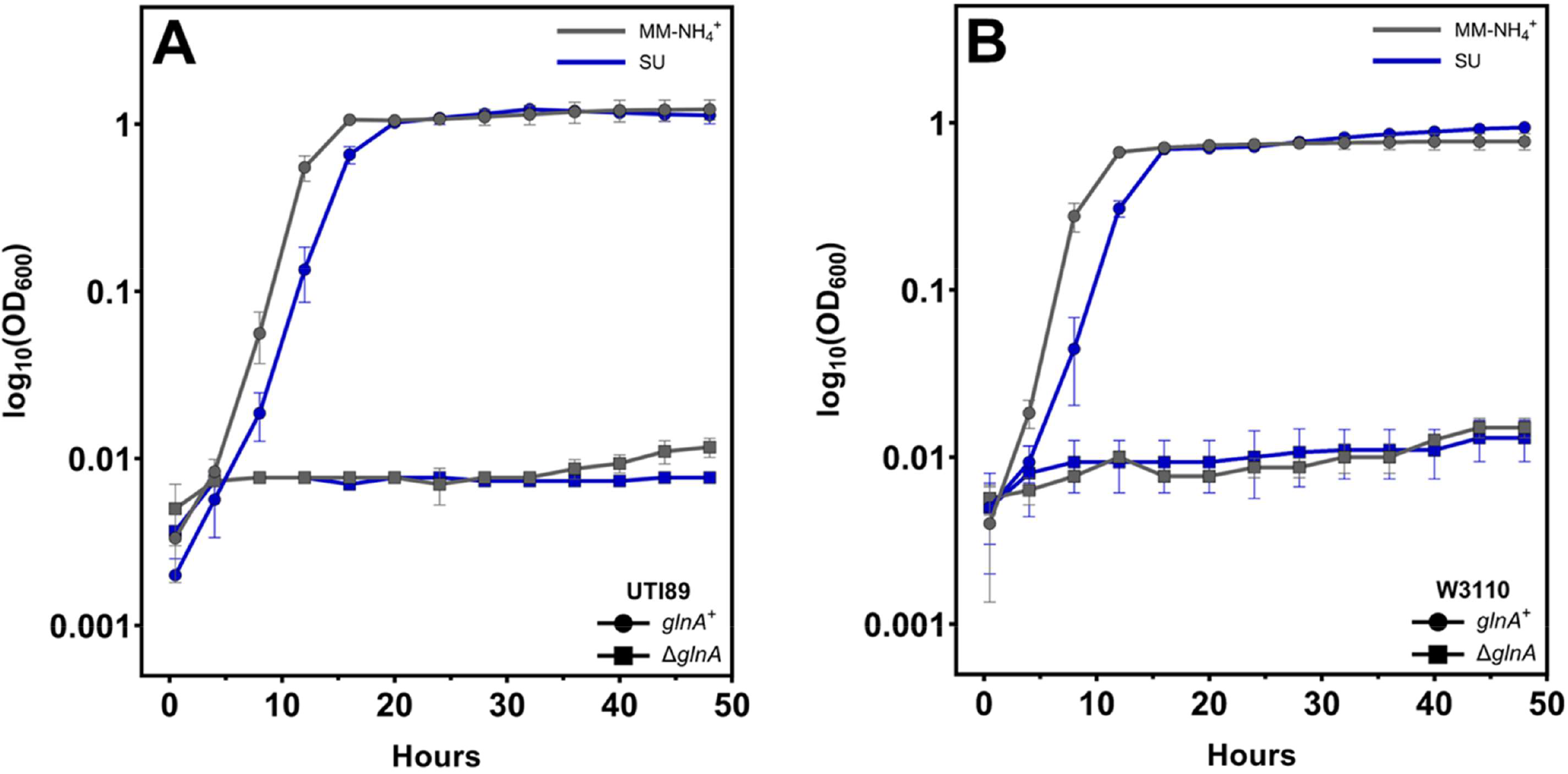
Growth of UTI89 and W3110 in minimal and basal SU medium. Growth curves of (A) UTI89 and (B) W3110 strains in the indicated media. The parental strains showed growth comparable in both media, while the Δ*glnA* mutants failed to grow. The curves are averages of three independent experiments and the error bars represent the standard deviations. Doubling times are provided in supplemental tables.

To more closely emulate urinary conditions, we supplemented SU medium with amino acids which total 5-7 mM in urine (13). SU supplementation with the 10 most abundant urinary amino acids (SU-10) did not affect UTI89 growth (Fig. 3A), but impaired W3110 growth (Fig. 3B). SU supplementation with the five most abundant urinary amino acids (SU-5) impaired UTI89 growth (Fig. 3A) and eliminated W3110 growth (Fig. 3B). Glutamine reversed the inhibitory effect (Fig. S2) which is consistent with the known inhibition of GS activity by the amino acids in SU-5 medium (14).

**Figure 3:**
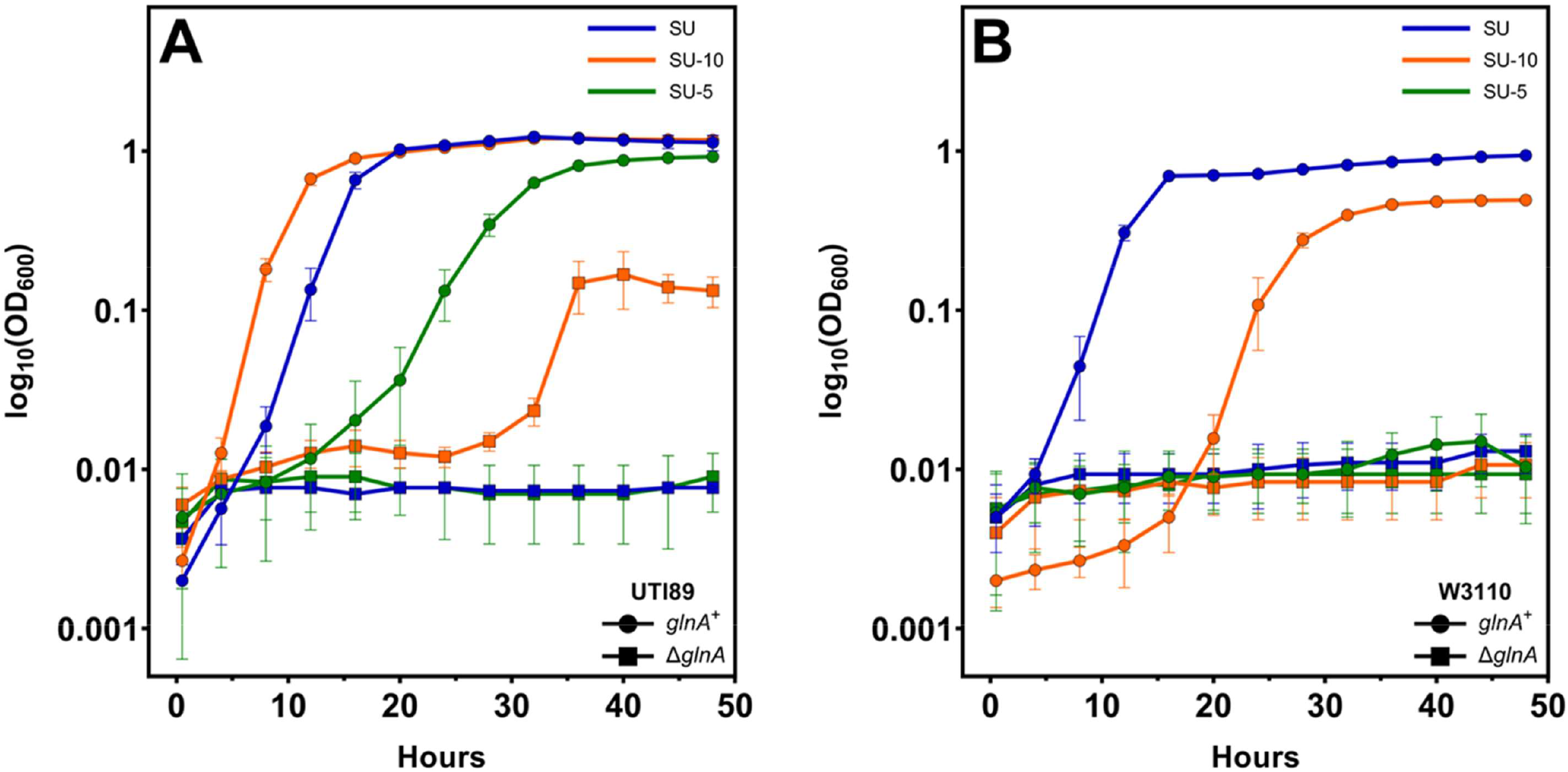
Growth of UTI89 and W3110 in SU supplemented with amino acids. Growth of **(A)** UTI89 and **(B)** W3110 in the indicated media. UTI89 was less sensitive to inhibitory amino acids compared to W3110 (green curves), and UTI89 Δ*glnA* showed marginal growth in SU-10 medium. The curves are averages of three independent experiments and the error bars represent the standard deviations. Doubling times are provided in supplemental tables.

Because SU-10 supports better growth than SU-5, we tested whether the GS substrate glutamate, which is present in SU-10, but not SU-5, could overcome the inhibitory effect of amino acids in SU-5. Glutamate improved growth in SU-5 for UTI89 (Fig. 4A), but not for W3110 (Fig. 4B). Additional glutamate in SU-10 improved growth for W3110 which shows that glutamate was not toxic (Urs, K. and Reitzer, L. unpublished observation). Tryptone or casamino acids also improved growth in SU-5 for both UTI89 and W3110, but the positive effect was only partial for W3110 and occurred after an extremely long lag phase (Figs. 4AB). In summary, urinary levels of several amino acids hinder growth and glutamate or glutamine overcomes the impairment which implies that the inhibitory amino acids block GS activity.

**Figure 4:**
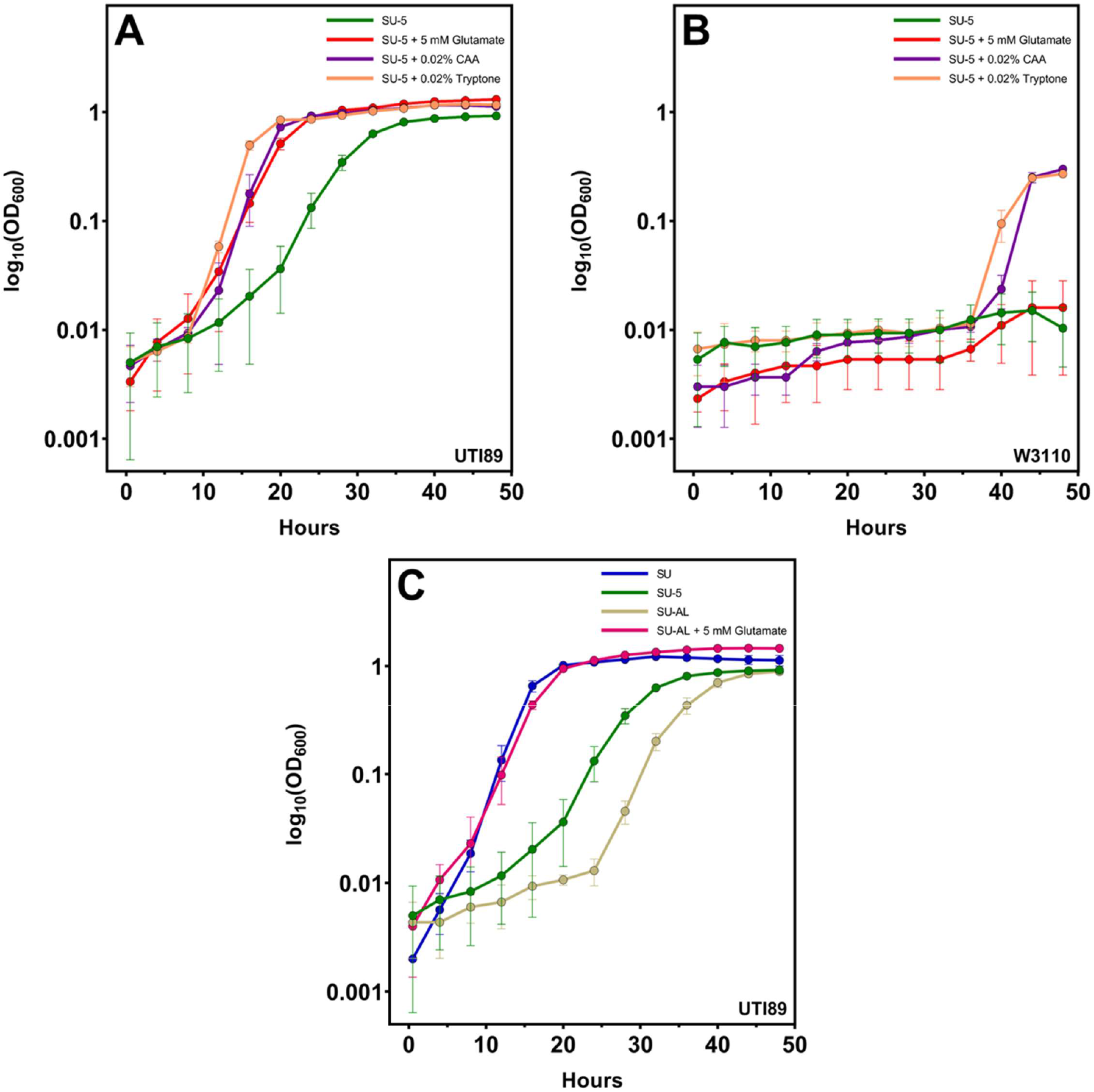
Glutamate or amino acid mixtures alleviated amino acid-induced growth inhibition in SU-5 medium or SU medium with the alanyl-leucine dipeptide. **(A)** Growth of WT UTI89 in SU-5 medium; **(B)** Growth of WT W3110 in SU-5 medium; and **(C)** Growth of WT UTI89 in SU-AL medium. W3110 did not grow in SU-AL medium. The curves are averages of three independent experiments and the error bars represent the standard deviations.

Urine consists of amino acids in free and peptide forms (13). Compared to free amino acids, dipeptides are transported rapidly into bacterial cells (15). A dipeptide containing an inhibitory amino acid should have the same effect as the five most abundant amino acids in urine. We supplemented SU with the alanyl-leucine (AL) dipeptide (SU-AL) which should generate the inhibitory amino acid alanine after transport and hydrolysis. UTI89 grown in SU-AL medium had a long lag phase which glutamate reduced (Fig. 4C). We conclude that the lag results from inhibition of GS. The AL dipeptide eliminated growth of W3110 in SU, which glutamate did not reverse (Urs, K. and Reitzer, L. unpublished observation). In summary, a single inhibitory amino acid can have the same effect as several inhibitory amino acids.

As a negative control for growth experiments, we grew Δ*glnA* derivatives of UTI89 and W3110, and as expected, Δ*glnA* mutants failed to grow in any media with one exception: UTI89Δ*glnA* displayed limited growth in SU-10 medium (Fig. 3A). We reconstructed UTI89Δ*glnA* several times, and the mutants always displayed the same phenotype. Furthermore, GS activity was undetectable in these mutants (Urs, K. and Reitzer, L. unpublished observation). An analysis of this phenotype is beyond the scope of this paper.

### Non-coordinate expression of glnA and the Ntr gene nac in urea-containing media

To test the effects of urine-like conditions on *glnA* expression, we grew strains carrying a plasmid with a transcriptional fusion of the *glnA* promoter region (*glnA* has two promoters) to the gene for the green fluorescent protein (*gfp*). The fusion properly responded to nitrogen availability in both strains for growth in minimal media: *glnA* expression was high in MM-alanine medium (nitrogen limiting) and low in MM-ammonia medium (nitrogen excess) (Figs. 5AB and quantified in Figs. 6AB). In ammonia-containing SU and SU-10 media, *glnA* expression was comparable to the activated level in MM-alanine in both strains (Figs. 5AB and 6AB). The presence of the 5 most abundant amino acids in urine or the alanyl-leucine dipeptide induced *glnA* expression about 3- to 4-fold higher than in SU medium (SU vs SU-5 and SU-AL), and glutamate reversed the increase (Figs. 6AB). In summary, *glnA* expression was induced by (a) amino acids that inhibit GS activity and (b) a component in SU medium.

**Figure 5:**
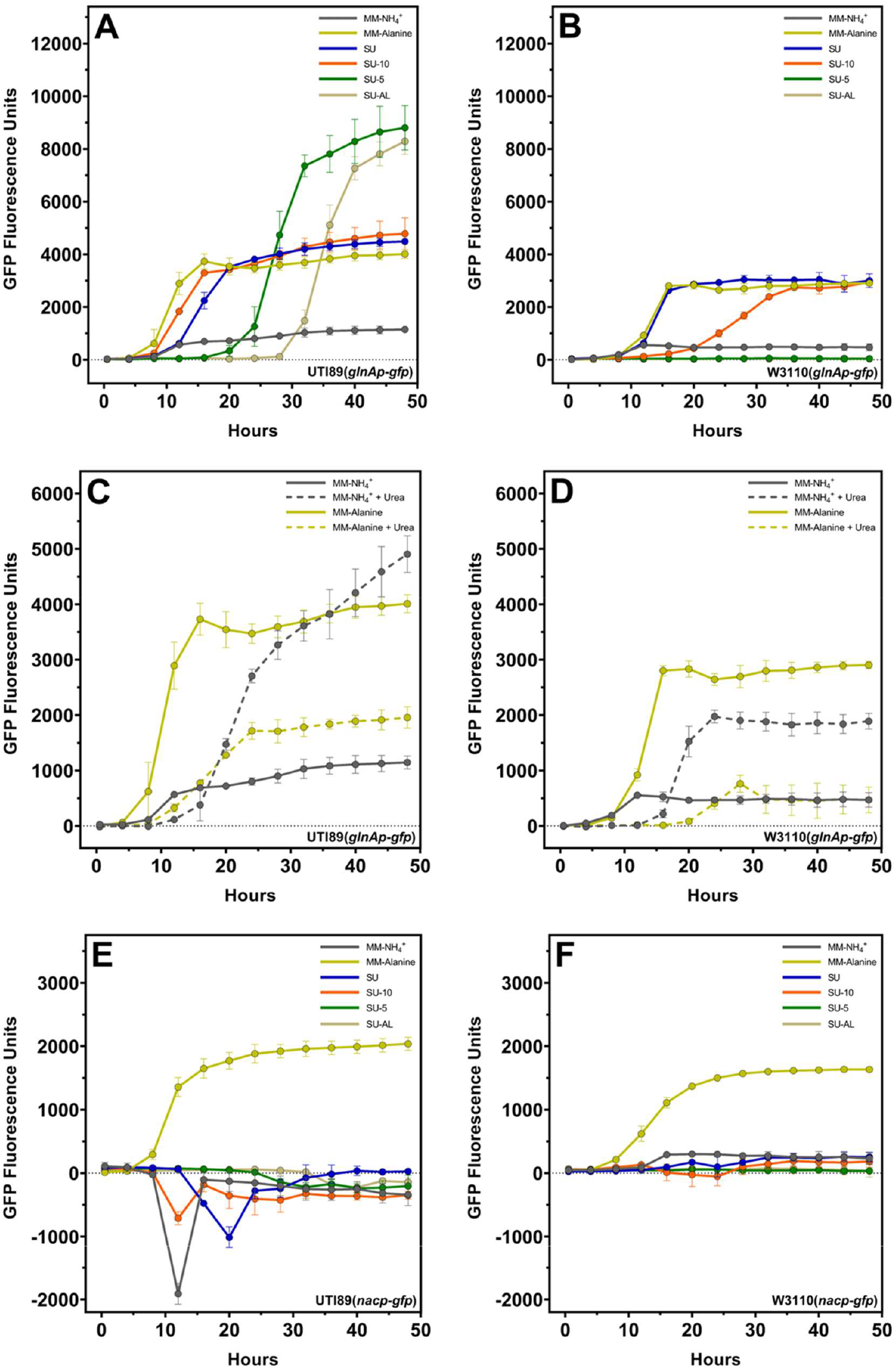
Effect of amino acids and urea on *glnA* and *nac* expression. *glnA* expression in the indicated media for **(A)** WT UTI89 and **(B)** WT W3110 – *glnA* expression in SU is comparable to expression levels in MM-alanine. Growth in SU-5 or SU-AL induces higher *glnA* expression for UTI89. *glnA* expression in nitrogen-limited (MM-alanine) and nitrogen-excess (MM-NH_4_^+^) minimal media with and without urea for **(C)** UTI89 and **(D)** W3110 – Urea induces *glnA* expression in MM-NH_4_^+^ but represses expression in MM-alanine. *nac* expression in the indicated media for **(E)** WT UTI89 and **(F)** WT W3110 – *nac* is not expressed during growth in SU in both strains. All curves represent average expression from three independent experiments measured concurrently with growth in respective media and error bars represent standard deviations.

**Figure 6:**
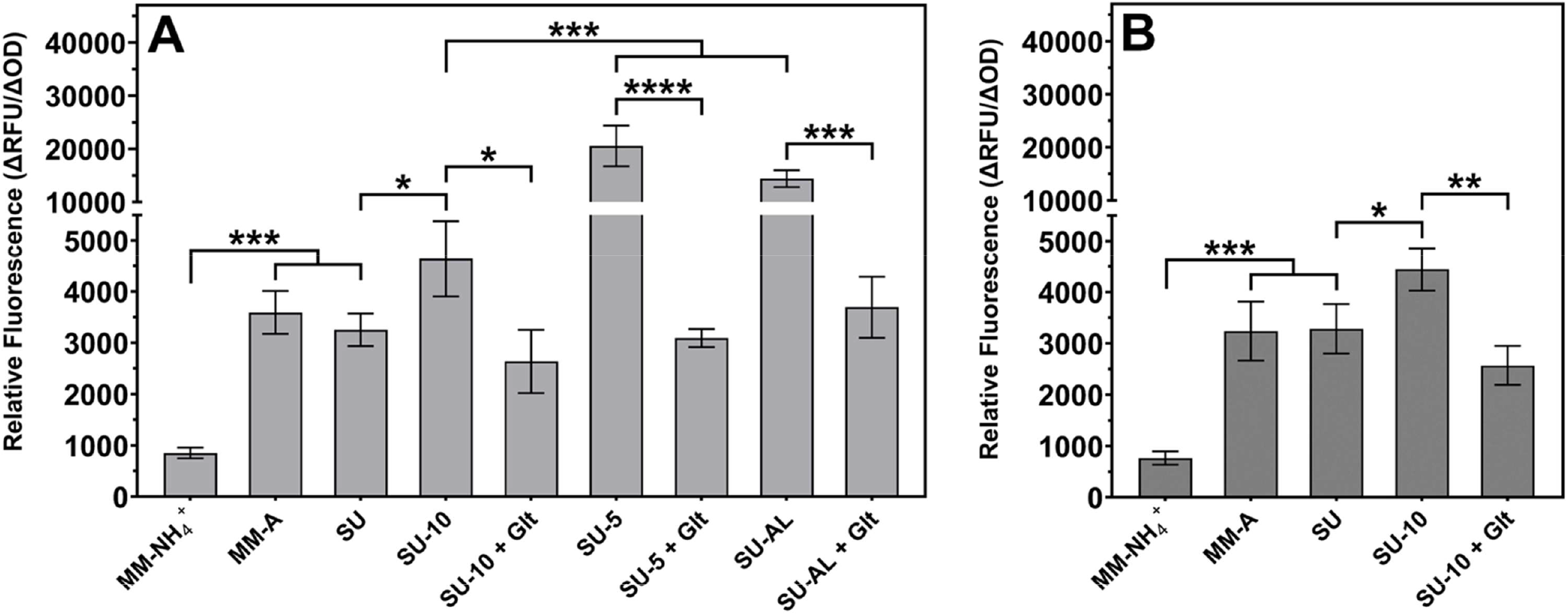
Relative transcriptional expression of *glnA* in different media. *glnA* expression in **(A)** WT UTI89 and **(B)** WT W3110. Because W3110 did not grow in SU-5 or SU-AL, there is no expression data. Significance was calculated using one-way ANOVA along with Dunnett’s multiple comparisons test; * p ≤ 0.05; ** p ≤ 0.01; *** p ≤ 0.001; **** p ≤ 0.0001. The data are averages of three independent experiments and the error bars represent the standard deviations.

Because SU medium does not contain amino acids that inhibit GS activity, induced *glnA* expression was unexpected. A major difference between SU and minimal media is urea. Addition of urea to MM-glucose-ammonia medium increased *glnA* expression 3-fold for both UTI89 and W3110, and decreased *glnA* expression in nitrogen-limited MM-glucose-alanine medium which does not contain ammonia (Figs. 5CD and quantified in Figs. 7AB) (similar results were observed for MM-glycerol medium ― Urs, K. and Reitzer, L. unpublished observation). Removal of urea from SU medium lowered *glnA* expression to the level for cells grown in MM-ammonia medium (Figs. 7AB), which would be expected for *glnA* expression in an ammonia-containing environment. The results of this section show that urea increased *glnA* expression in the presence of ammonia.

**Figure 7:**
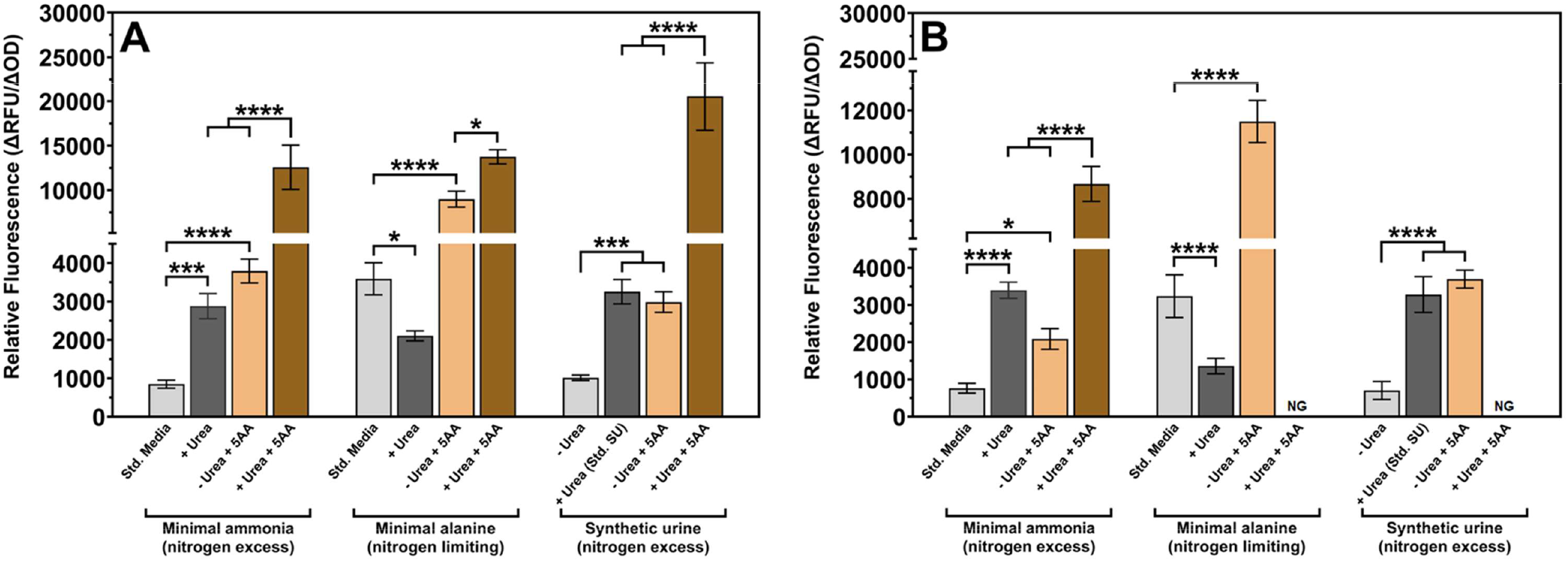
Relative transcriptional expression of *glnA* was altered by urea and the abundant urinary amino acids. **(A)** WT UTI89 and **(B)** WT W3110. NG means no growth. Significance was calculated using one-way ANOVA along with Dunnett’s multiple comparisons test; * p ≤ 0.05; ** p ≤ 0.01; *** p ≤ 0.001; **** p ≤ 0.0001. The data are averages of three independent experiments and the error bars represent the standard deviations.

*glnA* is part of the *glnALG* operon, which encodes the regulators of the Ntr response (14). To test if other Ntr genes are activated in parallel with *glnA*, we assayed the expression of *nac* (nitrogen assimilation control) which codes for a transcriptional regulator that activates a subset of Ntr genes (14, 16). Unlike *glnA* expression, *nac* expression was low in SU, SU-10, and SU-5 media (Figs. 5EF). To ensure that the fusion is properly regulated, we examined *nac* expression in minimal medium and, as expected, *nac* expression was high in both strains during growth in MM-alanine and low during growth in MM-ammonia (Figs. 5EF). In conclusion, *glnA* can be induced independent of other components of the Ntr response in synthetic urine.

### glnA expression from the Crp-dependent glnAp1 promoter in urea-containing media

Transcription of *glnA* is initiated at either the Crp-dependent *glnAp1* or the GlnG-dependent *glnAp2* promoter (Fig. 8) (17). Crp and GlnG are the only regulators of *glnA* expression, at least in the tested growth media, because Δ*glnG*Δ*crp* double mutants of UTI89 and W3110, like Δ*glnA* mutants, failed to grow (Fig. 9 and Figs. S3ABCD). The Δ*crp* mutants had an extended lag phase, often greater than 10 hours, for UTI89 and W3110 grown in SU (Fig. 9), and SU-10 and SU-5 (Fig. S3ABCD). Growth eventually occurred, but the final cell density was always less than the parental strains (Figs. 9 and S3). A *glnA*-containing plasmid eliminated the lag phase in the Δ*crp* strains (Figs. 9 and S3). The Δ*glnG* mutants did not have a lag phase and grew to approximately the same final cell density as the parental strains (Figs. 9 and S3). UTI89 Δ*glnG* grew slower than its parental strain, but W3110 Δ*glnG* grew as well as its parental strain (Figs. 9 and S3). Since GlnG-dependent transcription from *glnAp2* contributes to growth and *glnA* expression in a Δ*crp* mutant only after an extended lag period, Crp-dependent transcription from *glnAp1* is effectively required, even if not absolutely required, for growth in urea-containing media.

**Figure 8:**
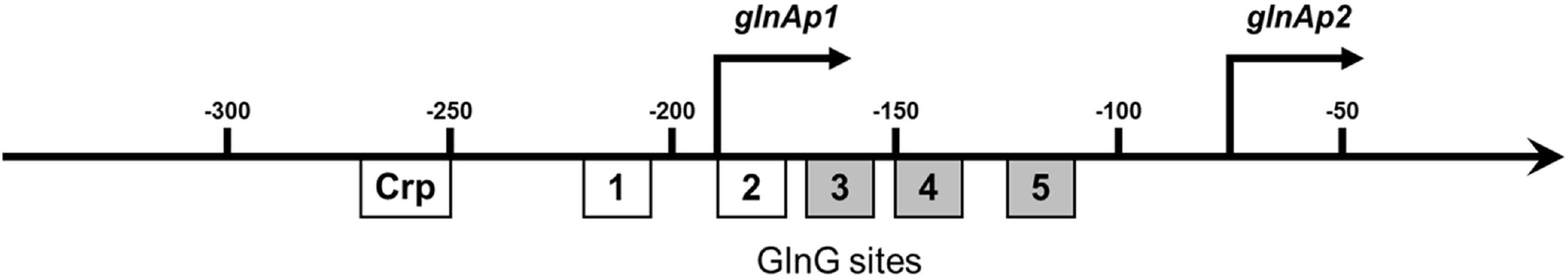
The *glnA* regulatory region. The binding sites are drawn to scale. GlnG sites 1 and 2 are strong sites required for activation from the *glnAp2* promoter, while sites 3-5 (shaded) appear to modulate activated expression (44). Crp-binding site(s) have not been experimentally determined for the *glnA* regulatory region. EcoCyc describes a Crp binding site from -133 to -120 which is between the strong GlnG binding sites and the RNA polymerase binding site (16).

**Figure 9:**
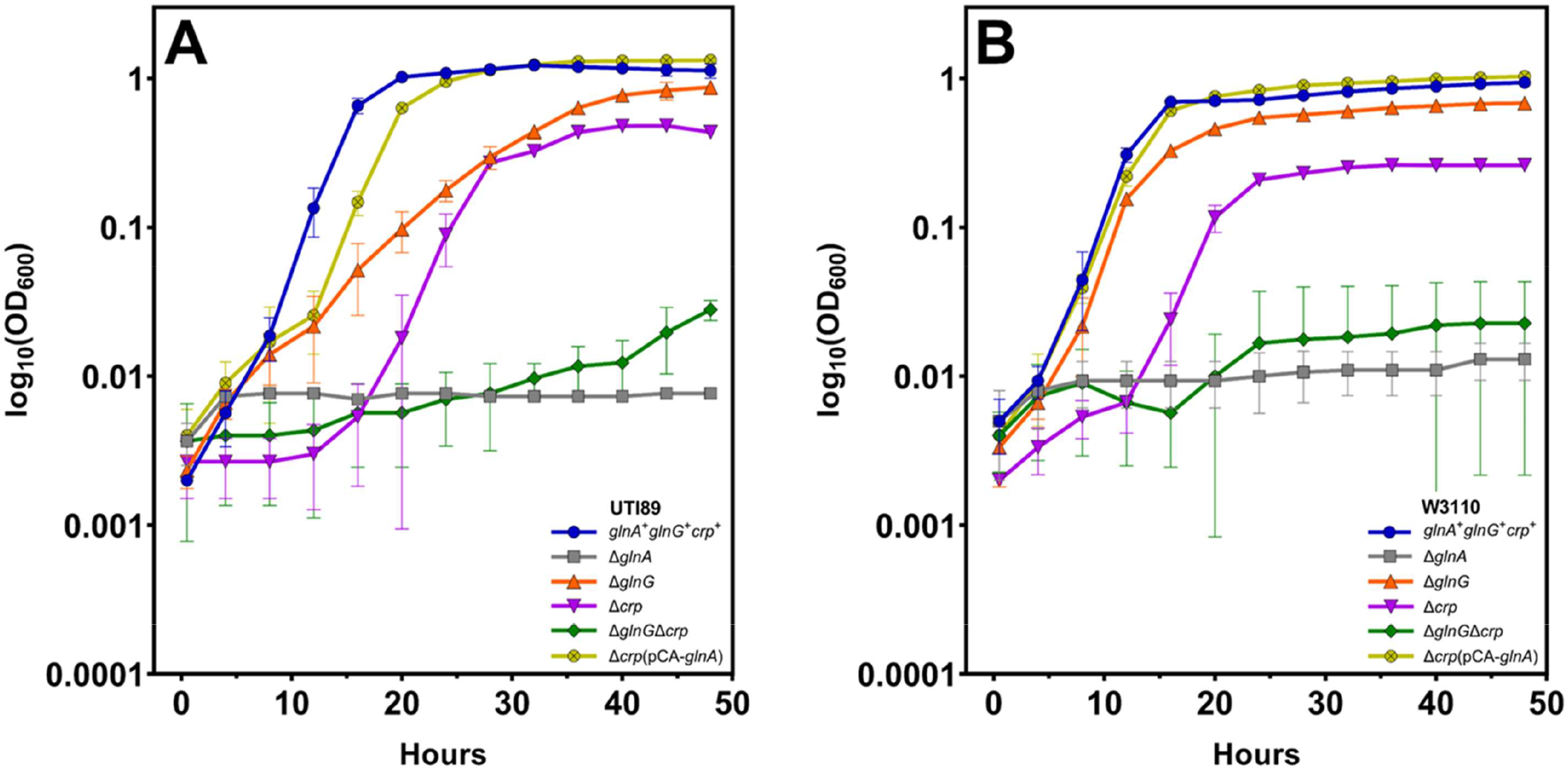
Growth phenotypes of Δ*glnG*, Δ*crp* and Δ*glnG*Δ*crp* mutants in SU medium. Growth of **(A)** UTI89 mutants and **(B)** W3110 mutants in SU medium. Growth is defective for Δ*crp* strains but not for Δ*glnG* strains. Overexpression of *glnA* in the Δ*crp* mutants restored growth back to WT levels in both strains. The curves are averages of three independent experiments and the error bars represent the standard deviations. Doubling times are provided in supplemental tables.

### Low glutamine synthetase activity and glnA translation during growth in urea-containing media

High *glnA* expression in SU medium is unexpected because of the high ammonia content. However, GS activity is very low and comparable to that in MM-ammonia (Figs. 10AB). GS activity in MM-alanine is significantly higher in comparison to the activity seen in any SU medium. The assay measures total GS activity regardless of the adenylylation state. To test the possibility that translational control could explain low enzymatic activity in cells with high *glnA* transcription, we constructed a plasmid containing a *glnA*-*gfp* translational fusion. Growth in urea-containing media resulted in expression from the translational fusion that was much lower than in nitrogen-limited minimal medium (Fig. 11), unlike expression from a *glnAp-gfp* transcriptional fusion (Fig. 6) for both UTI89 and W3110. In summary, elevated *glnA* transcription does not necessarily result in elevated GS synthesis.

**Figure 10:**
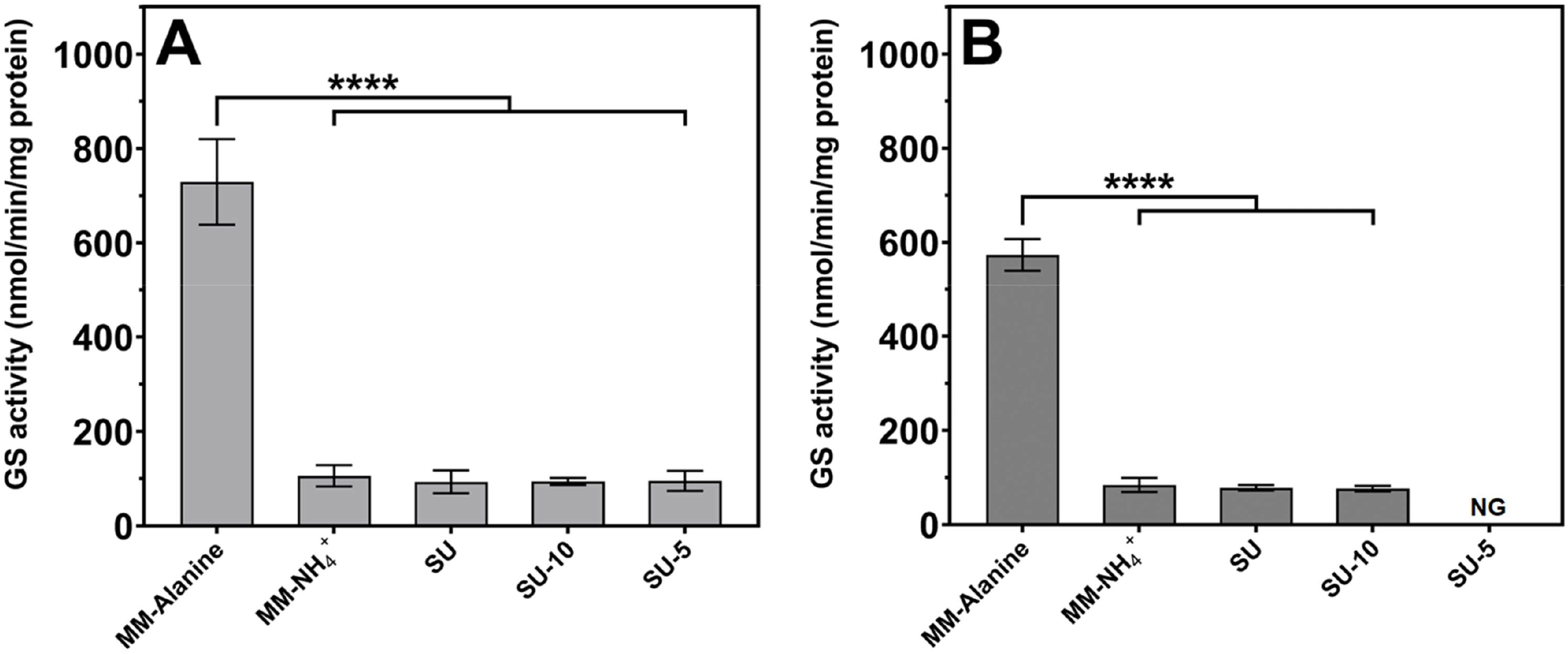
Glutamine synthetase activity during growth in various media. GS activity in **(A)** WT UTI89 and **(B)** WT W3110. NG means no growth. GS activity was not detectable in urea-containing media even though *glnA* was transcribed (see Figs. 5 and 6). Significance was calculated using one-way ANOVA along with Dunnett’s multiple comparisons test; **** p ≤ 0.0001. The data are averages of three independent experiments and the error bars represent the standard deviations.

**Figure 11:**
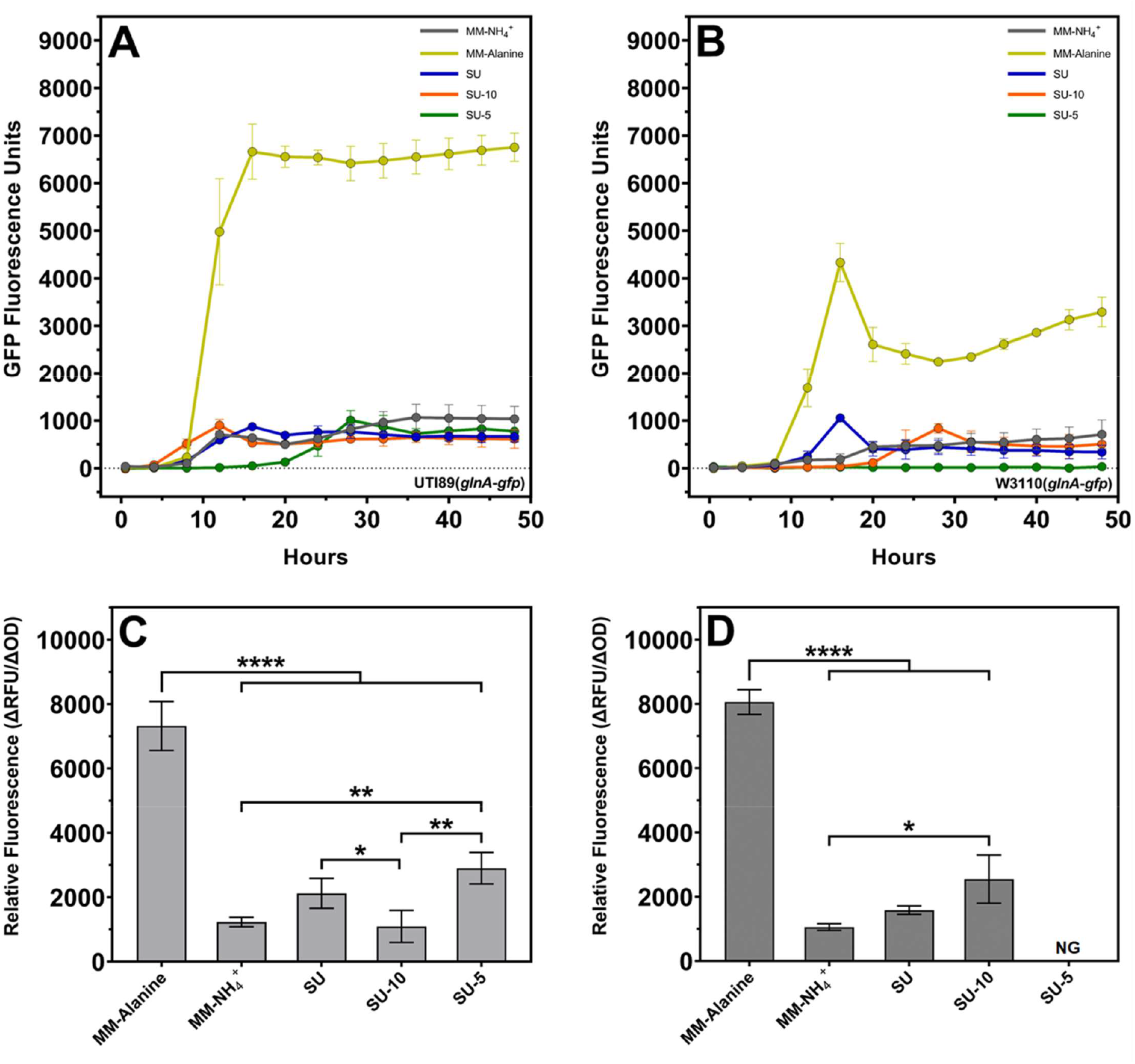
Translational expression of *glnA*. Expression during growth is shown for **(A)** WT UTI89 and **(B)** WT W3110 and quantified for **(C)** UTI89 and **(D)** W3110. NG means no growth. *glnA* mRNA was not translated during growth in urea-containing media even though *glnA* was transcribed (see Fig. 5). Significance was calculated using one-way ANOVA along with Dunnett’s multiple comparisons test; * p ≤ 0.05; ** p ≤ 0.01; **** p ≤ 0.0001. The data are averages of three independent experiments and the error bars represent the standard deviations.

## Discussion

*glnA* expression is unexpectedly high for *E. coli* grown in urine, despite a high ammonia concentration, which was interpreted to suggest that urine is nitrogen limited (11, 12). Our results describe activation of *glnA* expression by abundant components in urine — urea and several amino acids — in the presence of ammonia. Glutamate and glutamine reversed the activation by amino acids, but did not reverse the urea-dependent activation. These results imply that the abundant urinary amino acids inhibited GS activity, but that urea acts by a different mechanism. Urea-dependent *glnA* expression was initiated from the *glnAp1* promoter and did not result in enzymatically active GS or induction of the Ntr response. We conclude that *glnA* expression for cells grown in urine does not imply nitrogen limitation.

### Urea-dependent stimulation of *glnA* expression

The stimulatory effect of urea on *glnA* expression was observed for strains UTI89 and W3110 ― members of phylogenetic groups B2 and A, respectively ― which suggests that the urea effect is not strain specific. The mechanism of urea stimulation is unexpected because urea has been previously shown to inhibit expression from other cyclic-AMP-dependent promoters (18, 19). However, urea inhibited expression from the GlnG-dependent *glnAp2* promoter (lower *glnA* expression in MM-alanine medium with urea). Because transcription from *glnAp2* requires GlnG and supercoiled DNA (20, 21), one possibility is that a urea-dependent reduction of supercoiling in the *glnA* promoter region prevents one aspect of GlnG-dependent activation, such as binding to DNA, and consequent derepression from *glnAp1*. A second possibility is based on the known inhibition of *glnAp2* by Crp (22), if urea results in higher levels of cyclic-AMP, adenylate cyclase, or Crp. Regardless of mechanism, urea results in Crp-dependent *glnA* expression from *glnAp1*. Loss of Crp could result in a partial or complete glutamine auxotrophy for cells grown in a urinary environment, which could explain the attenuation of virulence of *crp* mutants in a mouse model (23). In this context, it should be noted that urinary urea in a mouse is six times higher than in humans (24), which could be a factor in mouse models of infection. Our results suggest that potential effects of urea on gene expression could contribute to virulence.

### Amino acid-dependent stimulation of *glnA* expression

Several amino acids have been known to inhibit GS activity for over 50 years (25, 26), and the mechanism of inhibition — binding to the glutamate site — is also known (27). However, the bladder is the first natural environment where the inhibitory amino acids are not only present at a high enough concentration to affect GS activity, but also *glnA* expression. This regulation is surprising because glutamate is by far the most abundant intracellular metabolite (28). However, alanine as the sole nitrogen source reduces intracellular glutamate (29), and the inhibitory amino acids may collectively have a similar effect. One of the inhibitory amino acids, alanine, has been previously shown to impair growth and alter gene expression in ammonia-containing minimal media (29, 30). Ikeda et al., citing unpublished results, argued that transcription in their ammonia-containing medium was initiated from the *glnAp2* promoter (29). If this conclusion is correct, then in the absence of urea the increase in *glnA* expression by our inhibitory amino acid mixture may result from transcription initiated from the *glnAp2* promoter (Fig 7). In the presence of urea, our results suggest that the inhibitory amino acids act via the *glnAp1* promoter because (a) the combined effects of urea and the amino acids were greater than urea alone (Fig. 7), urea impaired expression from the *glnAp2* promoter (Fig. 7, nitrogen limiting medium), and (c) loss of *crp* impaired growth more than loss of *glnG* in urea-containing media (Figs. 9 and S3). Although further experiments are required to more directly determine promoter usage during these conditions, our results clearly show that the inhibitory amino acids increase *glnA* expression regardless of media.

The responses to the inhibitory amino acids differed between UTI89 and W3110. SU-10 had no effect on UTI89 but inhibited growth of W3110, and SU-5 impaired growth of UTI89, and eliminated growth of W3110 (Fig 3). Our experience with these strains indicates that UTI89 and strains of phylogenetic group B2 in general grow faster in almost all media than W3110 and strains of phylogenetic group A (ref. (31) and Petter, A., Hogins, J. and Reitzer, L. unpublished observation). Of the 5 inhibitory amino acids, *E. coli* can degrade serine, glycine, alanine, and cysteine, but cannot degrade histidine (32, 33). We suggest that the differential responses may result from faster transport and degradation of the inhibitory amino acids by UTI89 that diminishes their intracellular concentration and minimizes their effect on *glnA* expression.

### Translational control of *glnA* expression

The unexpected and previously undescribed translational control of *glnA* was observed in urea-containing media (Fig. 10). Physical or chemical factors, such as urea which affects *glnA* expression but is not a sensor of nitrogen metabolism, may necessitate an additional layer of regulation. However, the factors that control this regulation are not apparent.

### Concluding remarks

A common assumption for analysis of pathogenic strains is that the regulation and physiology observed for growth in standard lab media for extensively passaged lab strains is the same as that for growth in natural environments. We provided evidence for previously undescribed aspects of nitrogen metabolism: urea-dependent stimulation of *glnA* expression, post-transcriptional control of *glnA* expression, and a curious observation that, superficially, is consistent with the possibility of *glnA*-independent glutamine synthesis, at least in UTI89. Study of *E. coli* strains other than laboratory strains in non-laboratory environments such as the bladder or bladder-like environments is likely to produce other surprises. For example, we showed *glnA* induction by the urinary components urea and several amino acids. GS and *glnA* would be a good therapeutic target for UTIs if urinary components were sufficiently high in all individuals to induce *glnA* expression. However, this is unlikely because of large variations in urinary components, changes in some components with age, and different levels of hydration that have the potential to dilute urinary components (13, 34). The initial interpretation that growth of *E. coli* in urine is nitrogen limited would imply that components of the Ntr response would be good therapeutic targets. Our results on the factors that affect *glnA* expression and their variations in the urine of individuals would suggest that *glnA* is not a good therapeutic target.

## Material and Methods

*Bacterial strains and plasmids*: All *E. coli* strains and plasmids used are listed in Table 1. Deletion alleles for Δ*glnG* and Δ*crp* were derived from the KEIO collection strains (35) and for Δ*glnA* from TH16 (36). Deletions were transferred to background strains by P1 transduction (37). Where necessary, the antibiotic resistance gene was removed using the pCP20 plasmid as described (38). Primers to check deletions are provided in Table S3. The low copy number plasmid pUA139 was obtained from the *E. coli* promoter collection, a library of fluorescent reporter strains carrying transcriptional fusions of GFP to different promoters in *E. coli* K12 strain MG1655 (39). Strains transformed with pUA139 and its derivatives carrying the upstream promoter regions of either *glnA* or *nac* were used to monitor gene expression during growth. The plasmid pCA24N encoding *glnA* under the control of IPTG-inducible promoters was derived from the ASKA collection (40) and used to assess the effect of *glnA* overexpression during growth in the deletion mutants.

**Table 1:**
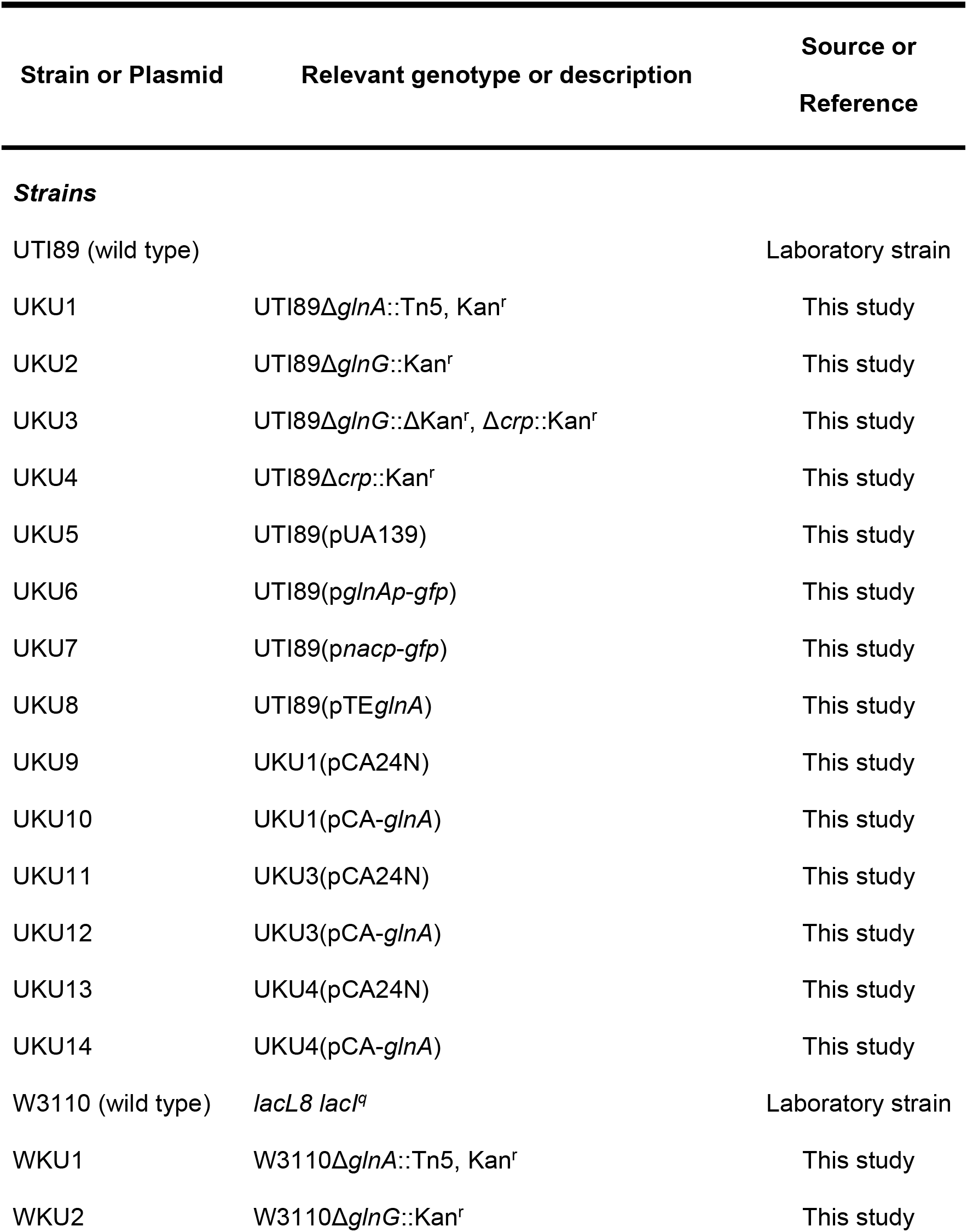
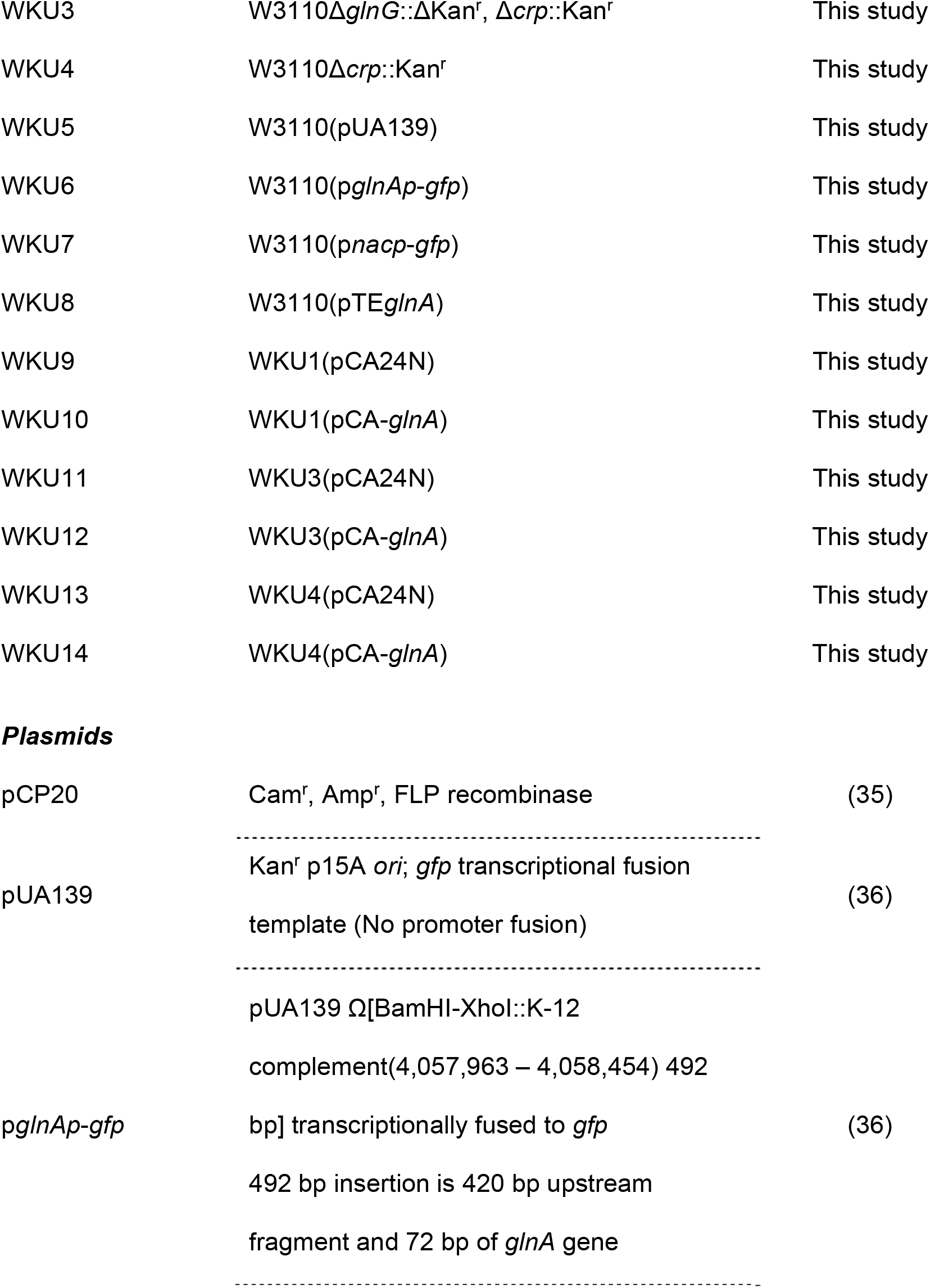
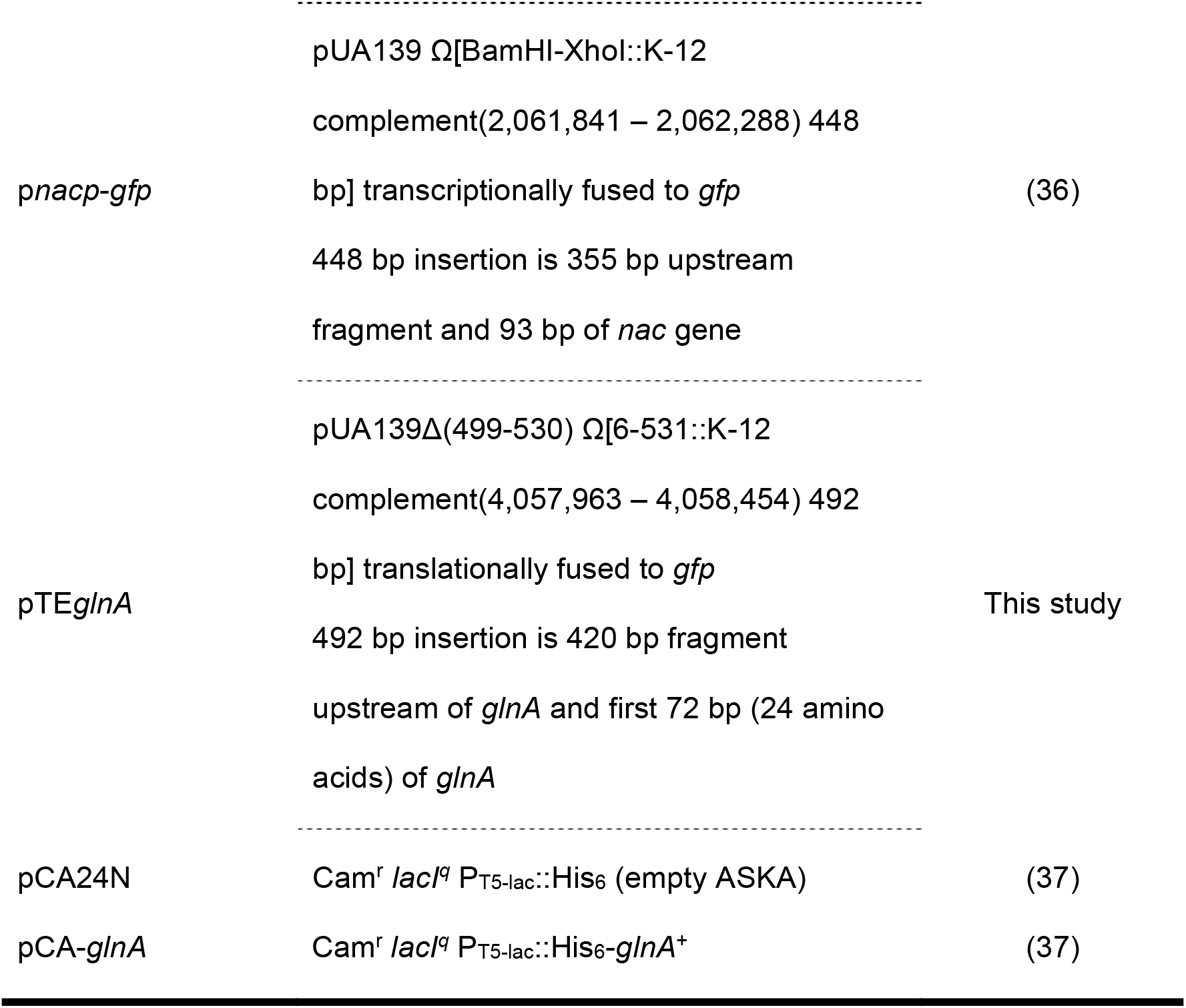
Strains and plasmids.

Plasmid pTE*glnA*, derived from pUA139, was constructed to monitor translational expression of *glnA* during growth. It carries the upstream promoter region of *glnA* along with an in-frame fusion of 72 base pairs (or 24 amino acids) of the N-terminal of GlnA to GFP. In brief, primers for separate *glnA* and *gfp* fragments were generated using the NEBuilder Assembly Tool (neb.com, New England Biolabs Inc., MA, USA). The fragments carry overlapping segments to one another and with the pUA139 vector. PCR amplified fragments were assembled with BamHI and SbfI linearized pUA139 using the NEBuilder HiFi DNA Assembly kit (New England Biolabs Inc., MA, USA) to generate pTE24G. Sequence verified plasmid was then used for transformation into strains. Primers used for amplification and sequencing are described in Table S3.

### Media and growth conditions

Strains were grown in either minimal medium (MM) containing ‘W’ salts (10.5 g/l K_2_HPO_4_, 4.5 g/l KH_2_PO_4_, and 0.05 g/l MgSO_4_ which was adjusted to pH 7.0) (41) or basal synthetic urine medium (SU) to assess growth, fluorescence and enzyme activity. The components for SU, SU-5, and SU-10 are listed in Table 2; the buffer was MES (50 mM) and the total mixture was adjusted to pH 6.0. Both media types contained 10 mM of a carbon source, which was glucose, unless otherwise indicated. MM was supplemented with 0.15% (w/v) nitrogen source ((NH_4_)_2_SO_4_ – nitrogen-rich or alanine – nitrogen-deficient). Commercial amino acid mixtures (tryptone or casamino acids), where indicated, were added at a concentration of 0.02% (w/v). To assess growth, starter cultures (12- to 14-hour incubation) from a single colony were grown either in MM-ammonia or SU. Starter cultures for the Δ*glnA*, Δ*crp* and Δ*glnG*Δ*crp* strains were supplemented with 5 mM glutamine. Cells were pelleted, washed with phosphate-buffer-saline (pH 7.4), resuspended in no-carbohydrate MM or SU and diluted to an OD_600_ of 0.2. Washed cells were inoculated at a ratio of 1:200 in a culture volume of 200 µl per well of a 96-well microtiter plate. Growth was assayed by measuring OD_600_ at intervals of 30 minutes for 48–72 hours on plate readers (Biotek Instruments Inc., VT, USA). Cultures were incubated at 37°C and at 237 cpm. For strains carrying the pUA139 plasmid or its derivatives, fluorescence was measured concurrently with growth at an excitation wavelength of 485 nm and emission recorded at 540 nm (42). IPTG at a final concentration of 0.2 mM was added to cultures carrying the pCA-*glnA* plasmid to control *glnA* expression. Strains carrying the empty pCA24N vector were used as controls to assess the effect of *glnA* expression during growth in the deletion mutants. Antibiotics, where necessary, were added at the following concentrations, kanamycin 25 µg/ml and chloramphenicol 7.5 µg/ml.

**Table 2:** Synthetic urine media components.

| **Component** | **Concentration in SU**<br>**(synthetic urine)** | **SU-5** | **SU-10** |
| --- | --- | --- | --- |
| Urea | 250 mM | as SU | as SU |
| NaCl | 100 mM | as SU | as SU |
| KCl | 40 mM | as SU | as SU |
| Na<sub>2</sub>HPO<sub>4</sub> | 10 mM | as SU | as SU |
| (NH<sub>4</sub>)<sub>2</sub>SO<sub>4</sub> | 10 mM | as SU | as SU |
| MgCl<sub>2</sub> | 3 mM | as SU | as SU |
| CaCl<sub>2</sub> | 3 mM | as SU | as SU |
| FeSO<sub>4</sub> | 0.005 mM | as SU | as SU |
| Glycine |  | 2.1 mM | as SU-5 |
| Histidine |  | 1.2 mM | as SU-5 |
| Cysteine |  | 0.6 mM | as SU-5 |
| Serine |  | 0.5 mM | as SU-5 |
| Alanine |  | 0.5 mM | as SU-5 |
| Leucine |  |  | 0.4 mM |
| Aspartate |  |  | 0.2 mM |
| Threonine |  |  | 0.2 mM |
| Methionine |  |  | 0.1 mM |
| Glutamate |  |  | 0.1 mM |

### Growth kinetics and relative fluorescence calculations

Data from each growth run were exported from the Gen5 software (Biotek Instruments Inc., VT, USA) into Microsoft excel and OD_600_ reads corrected for pathlength. Data from independent experiments were used to plot growth curves using GraphPad Prism (Ver. 9.2.1, GraphPad Software Inc., CA, USA). Specific growth rate (µ) of each culture was calculated using the formula, µ = ln(OD_t2_/OD_t1_)/(t2-t1), where OD_t2_ and OD_t1_ are OD_600_ during the exponential growth phase at time t2 and t1 respectively. Doubling time is calculated as, DT = ln(2)/µ. All time measurements are in hours. Final cell densities were determined as colony forming units (CFU) per milliliter. At the end of each growth assay, cultures were diluted and 20 µl spotted on Luria-Bertani (LB – 10 g tryptone, 10 g NaCl, 5 g yeast extract, pH 7.0) agar plates and incubated at 37°C for 16–18 hours before counting colonies for CFU calculations. For the fluorescence measurements, fluorescence reads from strains with an empty pUA139 vector (plasmid with no promoter region or truncated gene fused to GFP) were subtracted from the reads of strains carrying the p*glnAp*-*gfp*, p*nacp*-*gfp* and pTE*glnA* plasmids, under similar growth conditions, to correct for the background fluorescence signal at each time point. The corrected reads were used for calculating relative fluorescence, which is determined as the ratio of the rate of change of fluorescence to the rate of change of absorbance (OD_600_) during the exponential growth phase. All graphs were generated, and analyses performed using GraphPad Prism (Ver. 9.2.1, GraphPad Software Inc., CA, USA).

### Enzyme activity

Cells were harvested in the exponential phase during growth in microtiter plates by the addition of cetyltrimethylammonium bromide (CTAB) and MnCl_2_ to a final concentration of 100 µg/ml and 1 mM respectively, followed by shaking for 15 minutes. Cells were pelleted, resuspended in cold 2.5 mg/ml CTAB, pelleted again, resuspended in buffer (pH 7.27) containing 20 mM imidazole, 100 µg/ml CTAB and 0.3 mM MnCl_2_, stored on ice and used for the enzyme assays. Glutamine synthetase activity of cultures was determined by using 25-100 µl of the prepared cell suspension in a reaction mixture (pH 7.27) containing 135 mM imidazole, 80 µg/ml CTAB, 25 mM potassium arsenate, 20 mM hydroxylamine, 0.4 mM ADP, and 0.2 mM MnCl_2_. Glutamine at a final concentration of 20 mM was added to the reaction mixture incubated at 37°C. The reaction was stopped by adding 1 ml of a stop mix containing 0.2 M FeCl_3_, 0.12 M trichloroacetic acid and 0.25 N HCl. Cell debris were removed by centrifugation and the absorbance measured at 540 nm. Protein estimation was done by the Lowry method (43). Final enzyme activities are recorded as nanomoles of product formed per minute per milligram of protein. All graphs were generated, and analyses performed using GraphPad Prism (Ver. 9.2.1, GraphPad Software Inc., CA, USA).

## Acknowledgements

This work was supported by a UT Dallas Collaborative Biomedical Research Award grant program, but not by specific grants from any funding agency in the public, commercial, or not-for-profit sectors. The authors have no relevant affiliations or financial involvement with any organization or entity with a financial interest in or financial conflict with the subject matter or materials discussed.

## Figure Legends

**Table S1:**
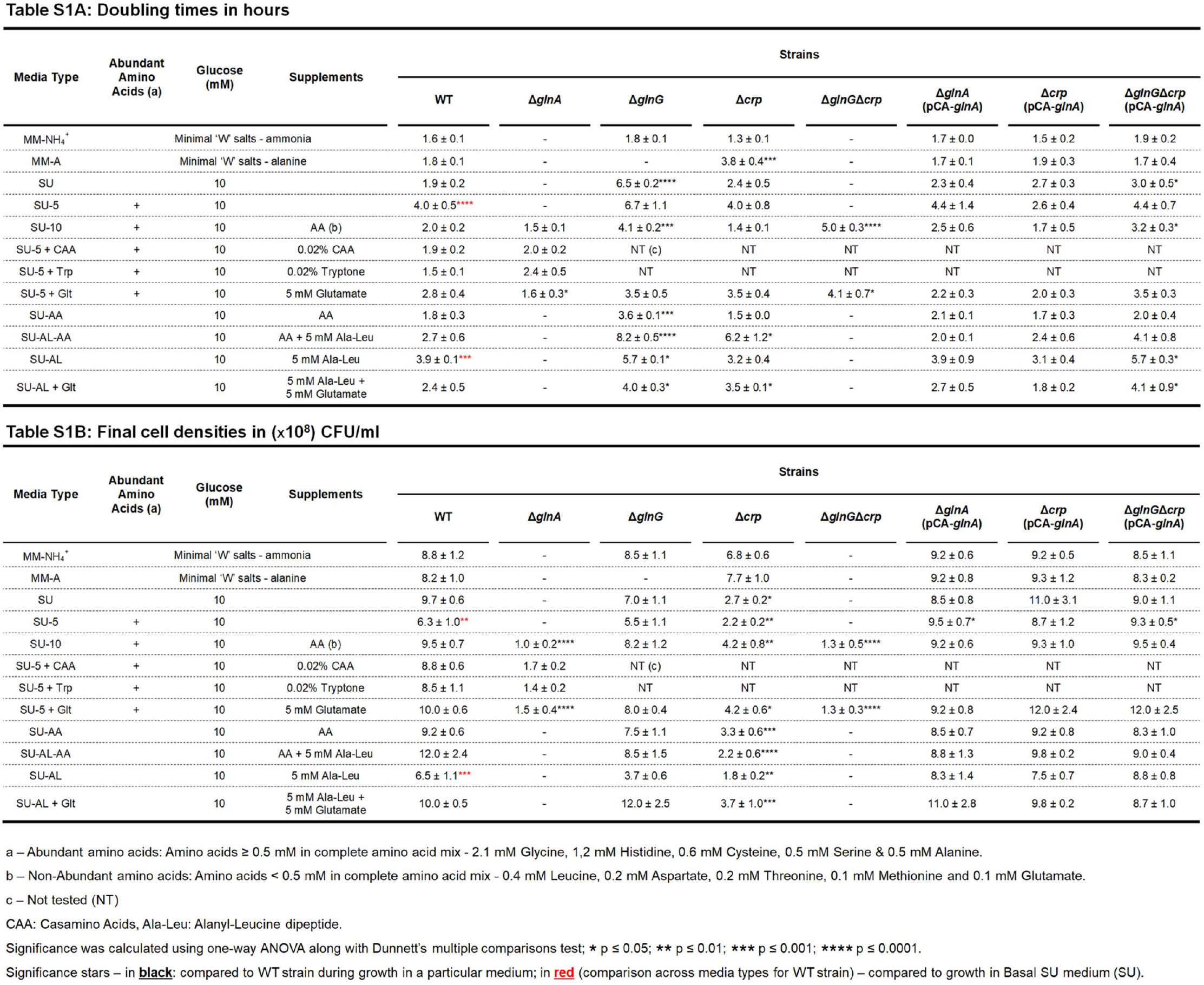
Growth kinetics of UTI89 Strains.

**Table S2:**
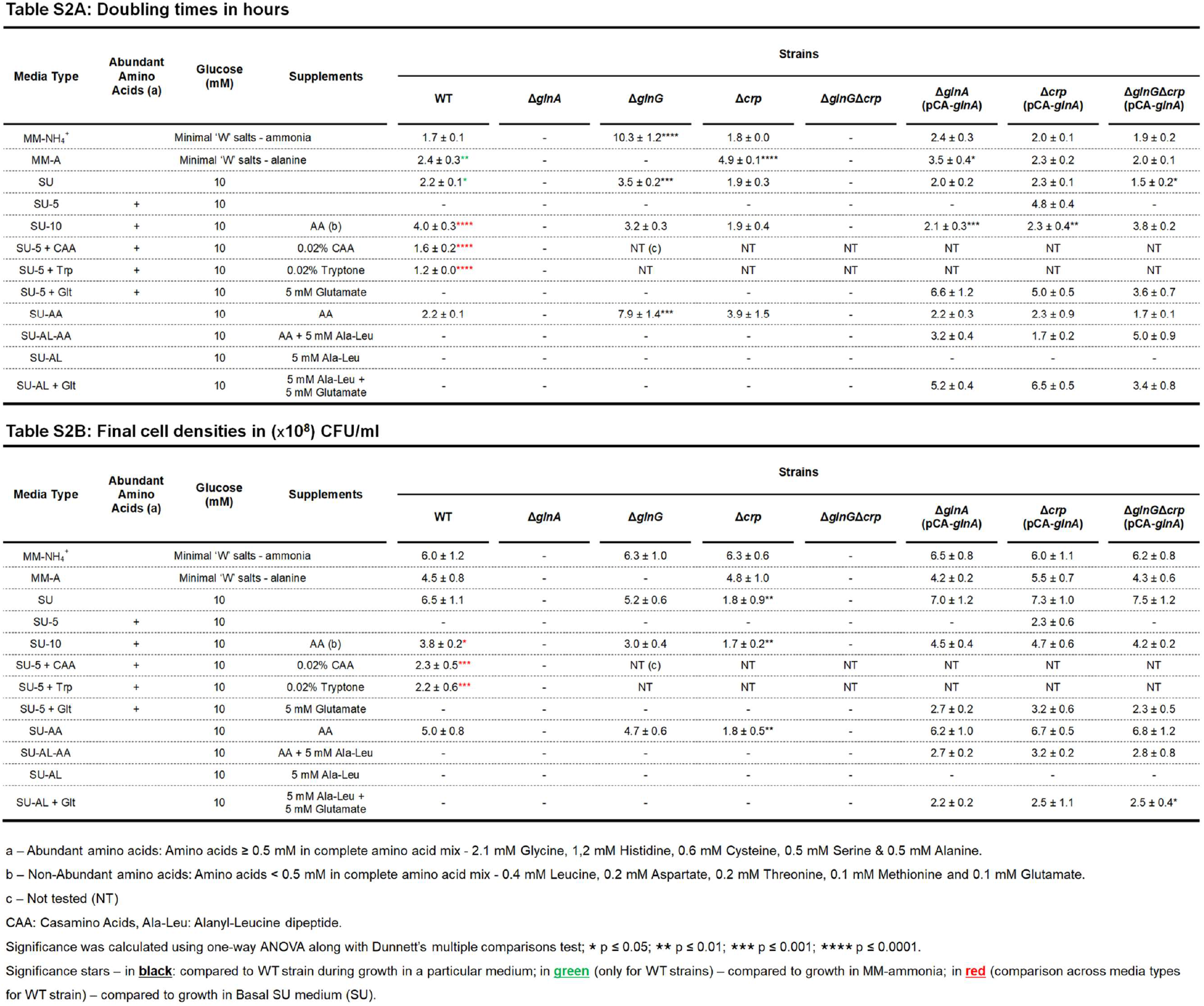
Growth kinetics of W3110 Strains.

**Table S3:**
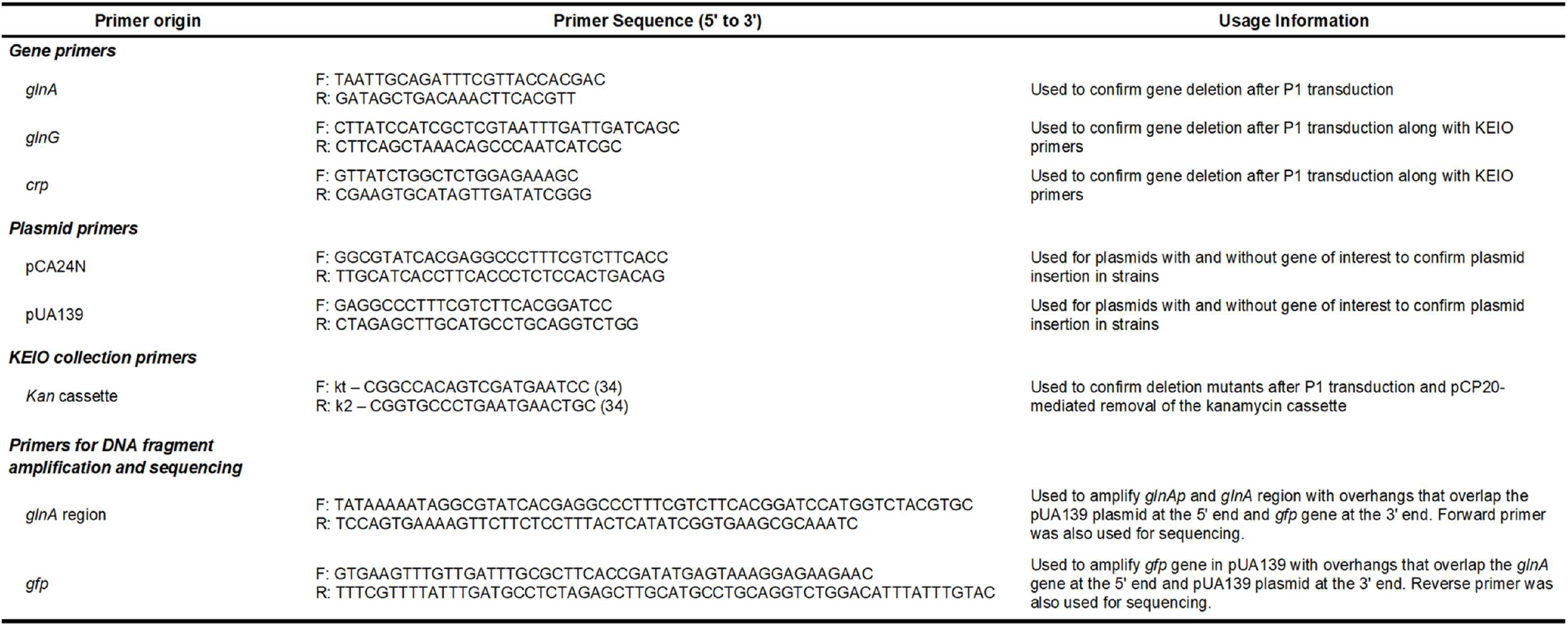
Primers used.

**Figure S1:**
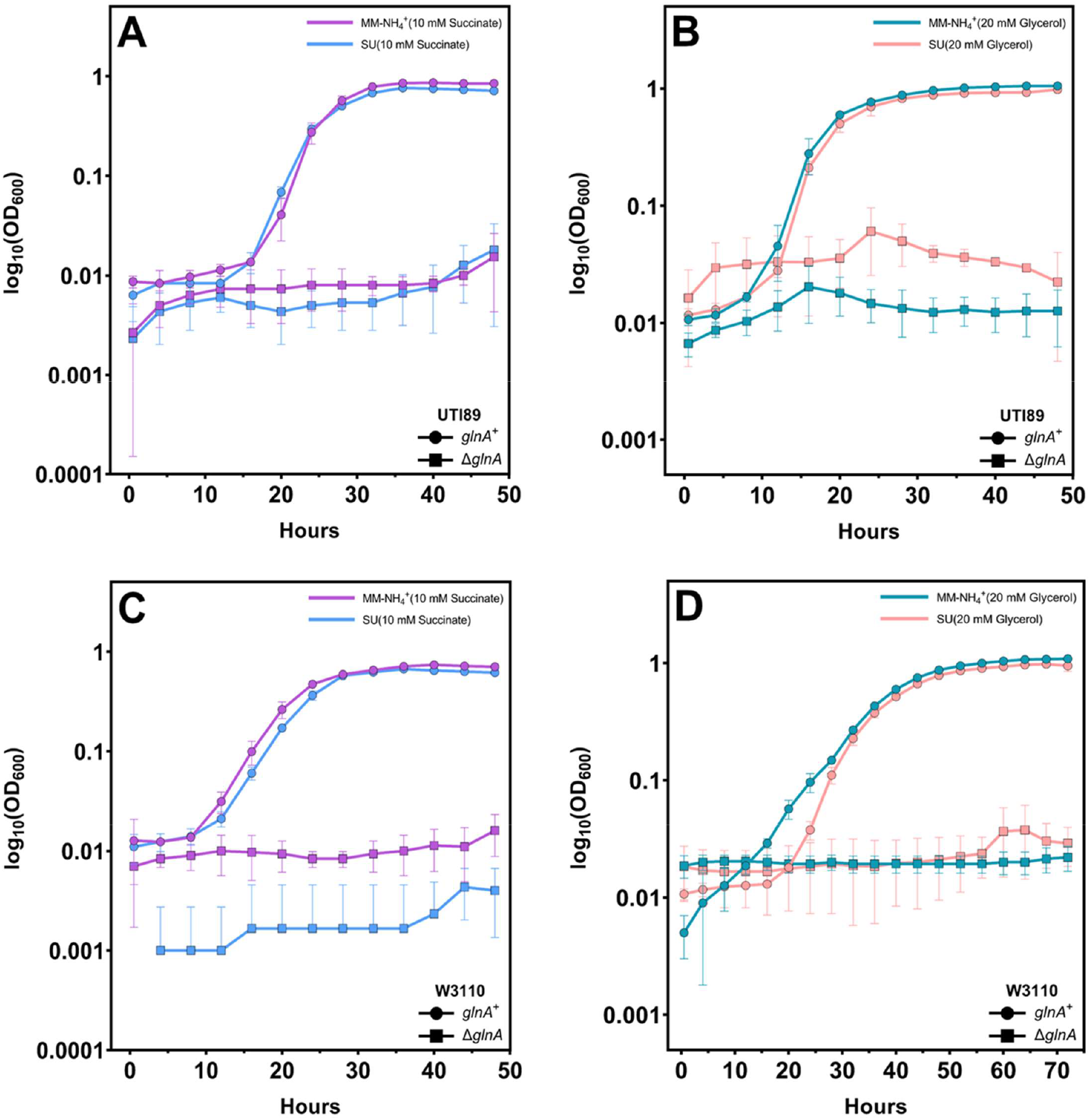
Growth of UTI89 and W3110 in Basal SU Medium: Growth of UTI89 strains in MM-NH_4_^+^ and SU medium with **(A)** 10 mM succinate or **(B)** 20 mM glycerol as the carbohydrate source. Growth of W3110 strains in MM-NH_4_^+^ and SU medium with **(C)** 10 mM succinate or **(D)** 20 mM glycerol as the carbohydrate source. The Δ*glnA* mutants for both UTI89 and W3110 show no viability in any medium. The curves are averages of three independent experiments and the error bars represent the standard deviations.

**Figure S2:**
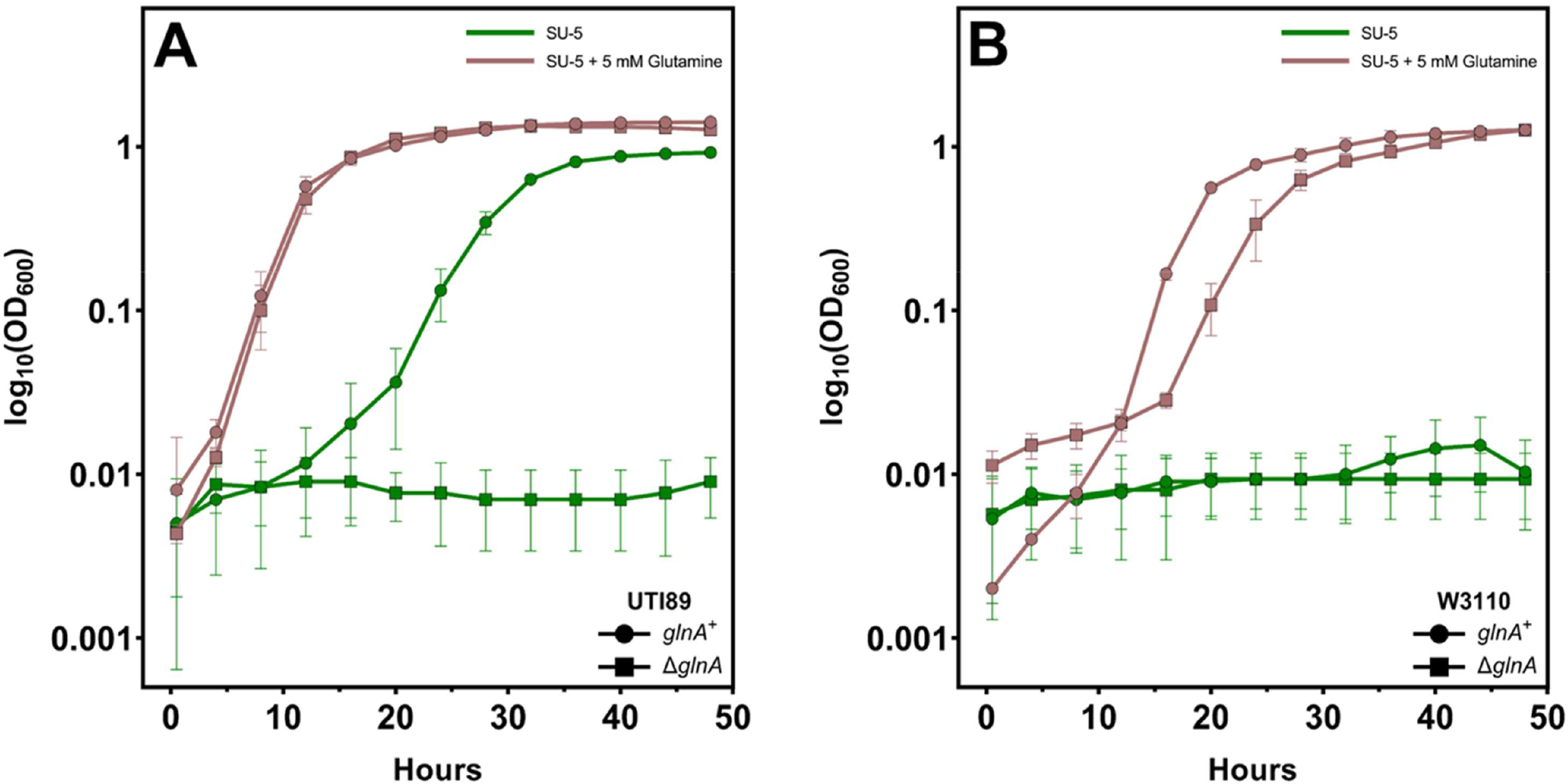
Growth of UTI89 and W3110 in SU-5 medium with glutamine supplementation: Growth in SU-5 medium with and without glutamine supplementation for **(A)** UTI89 and **(B)** W3110 strains. The WT strains recover growth upon glutamine addition in SU-5 medium. The Δ*glnA* mutants for both UTI89 and W3110 also show growth comparable to the WT strains with glutamine in the medium. The curves are averages of three independent experiments and the error bars represent the standard deviations.

**Figure S3:**
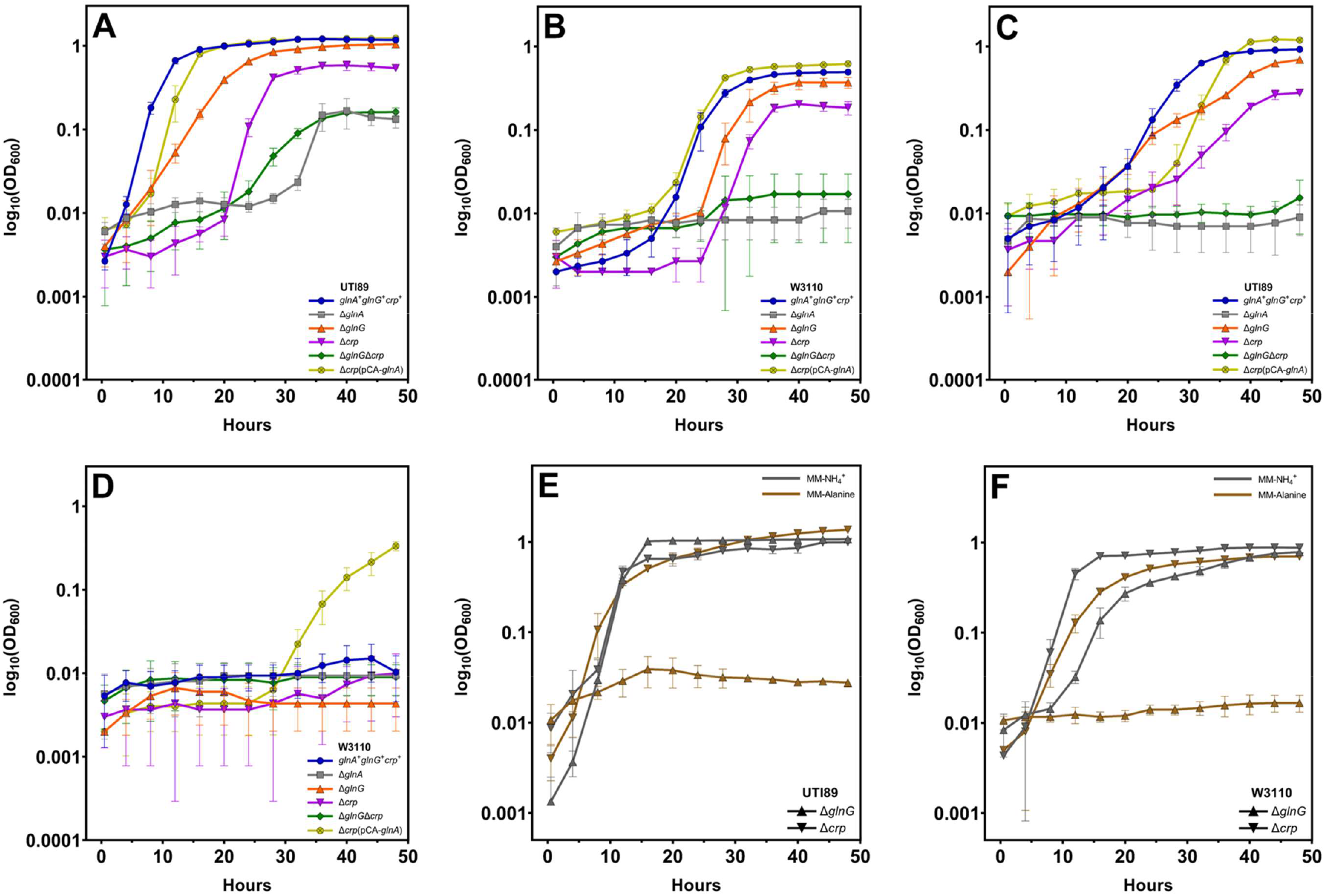
Growth of UTI89 and W3110 mutants: Growth of UTI89 and W3110 mutants in **(A) and (B)** SU-10 and in **(C) and (D)** SU-5 media - The Δ*glnG* mutants had phenotypes like the WT strains. The Δ*crp* mutants showed slower growth with significantly lower cell densities in both UTI89 and W3110. The Δ*glnG*Δ*crp* double mutants exhibited phenotypes like the Δ*glnA* mutant. Overexpression of *glnA* in the Δ*crp* mutants by an IPTG-inducible mechanism restored growth back to WT levels for UTI89. Overexpression of *glnA* in the W3110Δ*crp* mutant also allowed for marginal growth in SU-5. Growth of Δ*crp* and Δ*gInG* mutants in indicated media for **(E)** UTI89 and **(F)** W3110 - The Δ*crp* mutants grow as well as the WT strains in both MM-ammonia and MM-alanine media. The Δ*glnG* mutants for both UTI89 and W3110 showed no viability in a nitrogen-limited medium. The curves are averages of three independent experiments and the error bars represent the standard deviations.

## References

1. Ejrnaes K. 2011. Bacterial characteristics of importance for recurrent urinary tract infections caused by Escherichia coli. Dan Med Bull 58:B4187.

2. Foxman B. 2010. The epidemiology of urinary tract infection. Nat Rev Urol 7:653–660.

3. Rowe TA, Juthani-Mehta M. 2013. Urinary tract infection in older adults. Aging health. 9:doi:10.2217/ahe.13.38.

4. Flores-Mireles AL, Walker JN, Caparon M, Hultgren SJ. 2015. Urinary tract infections: epidemiology, mechanisms of infection and treatment options. Nat Rev Microbiol 13:269–284.

5. Forsyth VS, Armbruster CE, Smith SN, Pirani A, Springman AC, Walters MS, Nielubowicz GR, Himpsl SD, Snitkin ES, Mobley HLT. 2018. Rapid growth of uropathogenic Escherichia coli during human urinary tract infection. MBio. 9:doi:10.1128/mBio.00186-18.

6. Sintsova A, Frick-Cheng AE, Smith S, Pirani A, Subashchandrabose S, Snitkin ES, Mobley H. 2019. Genetically diverse uropathogenic Escherichia coli adopt a common transcriptional program in patients with UTIs. Elife. 8:doi:10.7554/eLife.49748.

7. Glover M, Moreira CG, Sperandio V, Zimmern P. 2014. Recurrent urinary tract infections in healthy and nonpregnant women. Urol Sci 25:1–8.

8. Selekman RE, Shapiro DJ, Boscardin J, Williams G, Craig JC, Brandström P, Pennesi M, Roussey-Kesler G, Hari P, Copp HL. 2018. Uropathogen resistance and antibiotic prophylaxis: a meta-analysis. Pediatrics. 142:doi:10.1542/peds.2018-0119.

9. Anonymous. 2021. Global antimicrobial resistance and use surveillance system (GLASS) report: 2021. World Health Organization, Geneva. https://apps.who.int/iris/handle/10665/341666.

10. Malik RD, Wu YR, Christie AL, Alhalabi F, Zimmern PE. 2018. Impact of allergy and resistance on antibiotic selection for recurrent urinary tract infections in older women. Urology 113:26–33.

11. Snyder JA, Haugen BJ, Buckles EL, Lockatell CV, Johnson DE, Donnenberg MS, Welch RA, Mobley HL. 2004. Transcriptome of uropathogenic Escherichia coli during urinary tract infection. Infect Immun 72:6373–6381.

12. Hagan EC, Lloyd AL, Rasko DA, Faerber GJ, Mobley HL. 2010. Escherichia coli global gene expression in urine from women with urinary tract infection. PLoS Pathog 6:e1001187.

13. Reitzer L, Zimmern P. 2020. Rapid growth and metabolism of uropathogenic Escherichia coli in relation to urine composition. Clin Microbiol Rev 33:e00101–00119.

14. Reitzer L. 2003. Nitrogen assimilation and global regulation in Escherichia coli. Annu Rev Microbiol 57:155–176.

15. Matthews DM, Payne JW. 1980. Transmembrane transport of small peptides, p 331–425. In Bronner F, Kleinzeller A (ed), Current Topics in Membranes and Transport, vol 14. Academic Press.

16. Karp PD, Ong WK, Paley S, Billington R, Caspi R, Fulcher C, Kothari A, Krummenacker M, Latendresse M, Midford PE, Subhraveti P, Gama-Castro S, Muniz-Rascado L, Bonavides-Martinez C, Santos-Zavaleta A, Mackie A, Collado-Vides J, Keseler IM, Paulsen I. 2018. The EcoCyc Database. EcoSal Plus. 8:doi:10.1128/ecosalplus.ESP-0006-2018.

17. Reitzer LJ, Magasanik B. 1985. Expression of glnA in Escherichia coli is regulated at tandem promoters. Proc Natl Acad Sci U S A 82:1979–1983.

18. Sanzey B, Ullmann A. 1976. Urea, a specific inhibitor of catabolite sensitive operons. Biochem Biophys Res Commun 71:1062–1068.

19. Sanzey B, Ullmann A. 1980. The effect of urea on catabolite sensitive operons in Escherichia coli K 12. Mol Gen Genet 178:611–616.

20. Claverie-Martin F, Magasanik B. 1992. Positive and negative effects of DNA bending on activation of transcription from a distant site. J Mol Biol 227:996–1008.

21. Hunt TP, Magasanik B. 1985. Transcription of glnA by purified Escherichia coli components: core RNA polymerase and the products of glnF, glnG, and glnL. Proc Natl Acad Sci U S A 82:8453–8457.

22. Tian ZX, Li QS, Buck M, Kolb A, Wang YP. 2001. The CRP-cAMP complex and downregulation of the glnAp2 promoter provides a novel regulatory linkage between carbon metabolism and nitrogen assimilation in Escherichia coli. Mol Microbiol 41:911–924.

23. Donovan GT, Norton JP, Bower JM, Mulvey MA. 2013. Adenylate cyclase and the cyclic AMP receptor protein modulate stress resistance and virulence capacity of uropathogenic Escherichia coli. Infect Immun 81:249–258.

24. Yang B, Bankir L. 2005. Urea and urine concentrating ability: new insights from studies in mice. Am J Physiol Renal Physiol 288:F881–896.

25. Woolfolk CA, Stadtman ER. 1967. Regulation of glutamine synthetase: III. Cumulative feedback inhibition of glutamine synthetase from Escherichia coli. Archives of Biochemistry and Biophysics 118:736–755.

26. Kingdon HS, Stadtman ER. 1967. Regulation of glutamine synthetase. X. Effect of growth conditions on the susceptibility of Escherichia coli glutamine synthetase to feedback inhibition. J Bacteriol 94:949–957.

27. Liaw SH, Pan C, Eisenberg D. 1993. Feedback inhibition of fully unadenylylated glutamine synthetase from Salmonella typhimurium by glycine, alanine, and serine. Proc Natl Acad Sci U S A 90:4996–5000.

28. Bennett BD, Kimball EH, Gao M, Osterhout R, Van Dien SJ, Rabinowitz JD. 2009. Absolute metabolite concentrations and implied enzyme active site occupancy in Escherichia coli. Nat Chem Biol 5:593–599.

29. Ikeda TP, Shauger AE, Kustu S. 1996. Salmonella typhimurium apparently perceives external nitrogen limitation as internal glutamine limitation. J Mol Biol 259:589–607.

30. Calvo JM, Matthews RG. 1994. The leucine-responsive regulatory protein, a global regulator of metabolism in Escherichia coli. Microbiol Rev 58:466–490.

31. Hogins J, Fan E, Seyan Z, Kusin S, Christie AL, Zimmern PE, Reitzer L. 2022. Bacterial growth of uropathogenic Escherichia coli in pooled urine is much higher than predicted from the average growth in individual urine samples. Microbiol Spectr. e0201622. doi:10.1128/spectrum.02016-22.

32. Reitzer L. 2005. Catabolism of amino acids and related compounds. EcoSal Plus. 1:doi:10.1128/ecosalplus.3.4.7.

33. Loddeke M, Schneider B, Oguri T, Mehta I, Xuan Z, Reitzer L. 2017. Anaerobic cysteine degradation and potential metabolic coordination in Salmonella enterica and Escherichia coli. J Bacteriol 199:e00117–00117.

34. Moriguchi J, Ezaki T, Tsukahara T, Fukui Y, Ukai H, Okamoto S, Shimbo S, Sakurai H, Ikeda M. 2005. Decreases in urine specific gravity and urinary creatinine in elderly women. Int Arch Occup Environ Health 78:438–445.

35. Baba T, Ara T, Hasegawa M, Takai Y, Okumura Y, Baba M, Datsenko KA, Tomita M, Wanner BL, Mori H. 2006. Construction of Escherichia coli K-12 in-frame, single-gene knockout mutants: the Keio collection. Mol Syst Biol 2:2006 0008.

36. Reitzer LJ, Magasanik B. 1986. Transcription of glnA in E. coli is stimulated by activator bound to sites far from the promoter. Cell 45:785–792.

37. Miller JH. 1972. Experiments in Molecular Genetics. Cold Spring Harbor Laboratory, Cold Spring Harbor, NY.

38. Datsenko KA, Wanner BL. 2000. One-step inactivation of chromosomal genes in Escherichia coli K-12 using PCR products. Proc Natl Acad Sci U S A 97:6640–6645.

39. Zaslaver A, Bren A, Ronen M, Itzkovitz S, Kikoin I, Shavit S, Liebermeister W, Surette MG, Alon U. 2006. A comprehensive library of fluorescent transcriptional reporters for Escherichia coli. Nat Methods 3:623–628.

40. Kitagawa M, Ara T, Arifuzzaman M, Ioka-Nakamichi T, Inamoto E, Toyonaga H, Mori H. 2005. Complete set of ORF clones of Escherichia coli ASKA library (a complete set of E. coli K-12 ORF archive): unique resources for biological research. DNA Res 12:291–299.

41. Smith GR, Halpern YS, Magasanik B. 1971. Genetic and metabolic control of enzymes responsible for histidine degradation in Salmonella typhimurium. 4-imidazolone-5-propionate amidohydrolase and N-formimino-L-glutamate formiminohydrolase. J Biol Chem 246:3320–3329.

42. Cormack BP, Valdivia RH, Falkow S. 1996. FACS-optimized mutants of the green fluorescent protein (GFP). Gene 173:33–38.

43. Lowry OH, Rosebrough NJ, Farr AL, Randall RJ. 1951. Protein measurement with the Folin phenol reagent. J Biol Chem 193:265–275.

44. Atkinson MR, Pattaramanon N, Ninfa AJ. 2002. Governor of the glnAp2 promoter of Escherichia coli. Mol Microbiol 46:1247–1257.

